# Copper drives remodeling of metabolic state and progression of clear cell renal cell carcinoma

**DOI:** 10.1101/2024.01.16.575895

**Authors:** Megan E. Bischoff, Behrouz Shamsaei, Juechen Yang, Dina Secic, Bhargav Vemuri, Julie A. Reisz, Angelo D’Alessandro, Caterina Bartolacci, Rafal Adamczak, Lucas Schmidt, Jiang Wang, Amelia Martines, Jacek Biesiada, Katherine E. Vest, Pier P. Scaglioni, David R. Plas, Krushna C. Patra, Shuchi Gulati, Julio A. Landero Figueroa, Jarek Meller, J. Tom Cunningham, Maria F. Czyzyk-Krzeska

**Affiliations:** Department of Cancer Biology, University of Cincinnati College of Medicine, Cincinnati, OH 45267, USA; Department of Biomedical Informatics, University of Cincinnati College of Medicine, Cincinnati, OH 45267, USA; Division of Biostatistics and Bioinformatics, Department of Environmental and Public Health Sciences University of Cincinnati College of Medicine, Cincinnati, OH 45267, USA; Division of Biomedical Informatics, Cincinnati Children’s Hospital Medical Center, Cincinnati, OH 45229, USA; Department of Biochemistry and Molecular Genetics, University of Colorado School of Medicine, Aurora, CO 80045, USA; Division of Hematology and Oncology, Department of Internal Medicine University of Cincinnati College of Medicine, Cincinnati, OH 45267, USA; Institute of Engineering and Technology, Faculty of Physics, Astronomy and Informatics, Nicolaus Copernicus University, Torun, 87-100, Poland; Trace Elements Group, Department of Environmental Medicine and Public Health at Icahn school of Medicine at Mount Sinai, NY 10029, USA; Department of Pathology and Laboratory Medicine, University of Cincinnati College of Medicine, Cincinnati OH 45219, USA; Division of Oncology and Hematology, Department of Internal Medicine, University of California Davis Comprehensive Cancer Center, Sacramento, CA 95817, USA; Department of Molecular and Cellular Biosciences, University of Cincinnati College of Medicine, Cincinnati, OH 45219, USA; Department of Computer Science, University of Cincinnati College of Engineering and Applied Sciences, Cincinnati, OH 45221, USA; Veteran Affairs Medical Center, Department of Veterans Affairs, Cincinnati, OH 45220, USA; Department of Pharmacology and System Biology, University of Cincinnati College of Medicine, Cincinnati, OH 45267, USA

## Abstract

Copper (Cu) is an essential trace element required for mitochondrial respiration. Late-stage clear cell renal cell carcinoma (ccRCC) accumulates Cu and allocates it to mitochondrial cytochrome c oxidase. We show that Cu drives coordinated metabolic remodeling of bioenergy, biosynthesis and redox homeostasis, promoting tumor growth and progression of ccRCC. Specifically, Cu induces TCA cycle-dependent oxidation of glucose and its utilization for glutathione biosynthesis to protect against H_2_O_2_ generated during mitochondrial respiration, therefore coordinating bioenergy production with redox protection. scRNA-seq determined that ccRCC progression involves increased expression of subunits of respiratory complexes, genes in glutathione and Cu metabolism, and NRF2 targets, alongside a decrease in HIF activity, a hallmark of ccRCC. Spatial transcriptomics identified that proliferating cancer cells are embedded in clusters of cells with oxidative metabolism supporting effects of metabolic states on ccRCC progression. Our work establishes novel vulnerabilities with potential for therapeutic interventions in ccRCC.

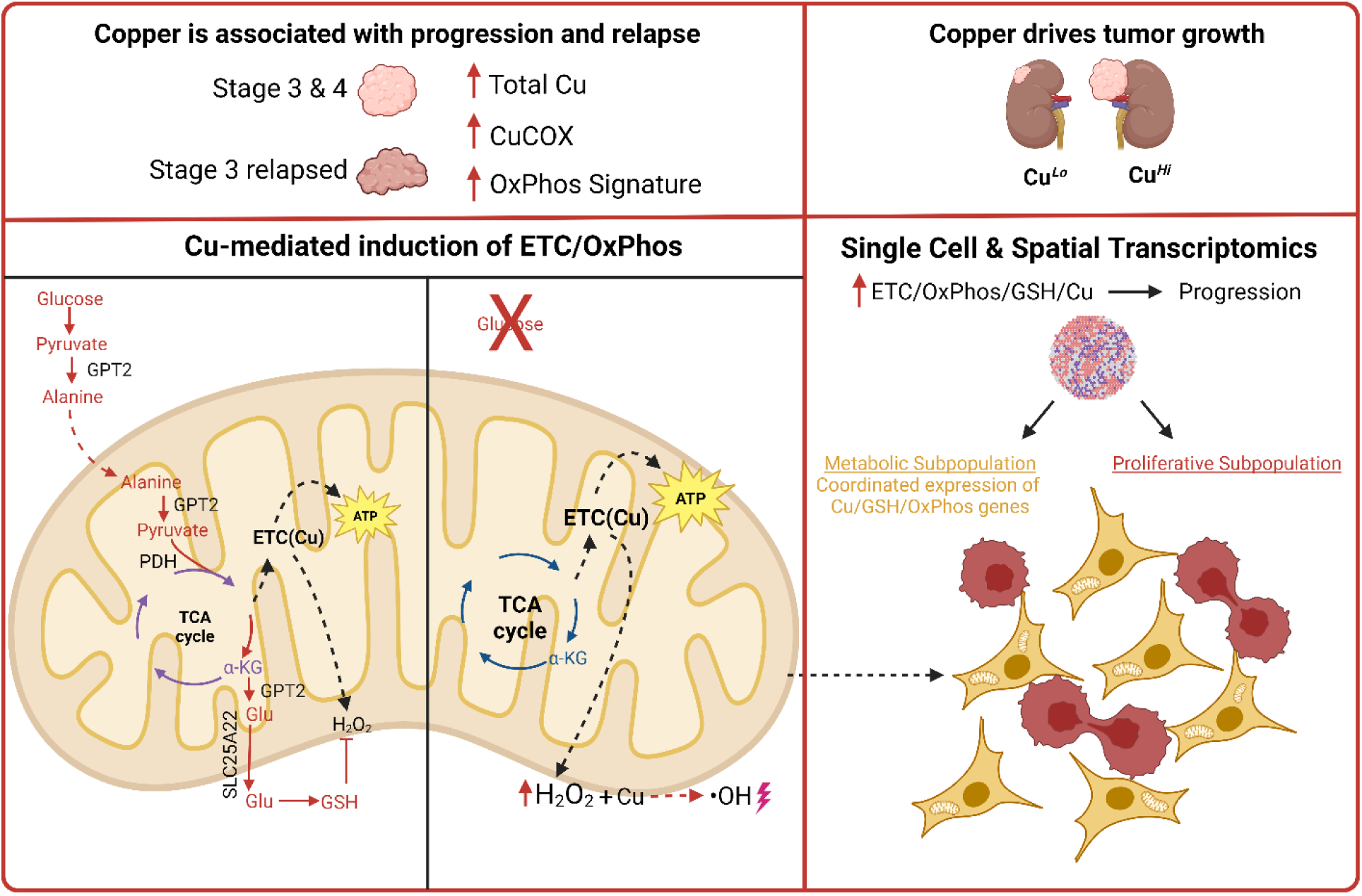

- Accumulation of copper is associated with progression and relapse of ccRCC and drives tumor growth.
- Cu accumulation and allocation to cytochrome c oxidase (CuCOX) remodels metabolism coupling energy production and nucleotide biosynthesis with maintenance of redox homeostasis.
- Cu induces oxidative phosphorylation via alterations in the mitochondrial proteome and lipidome necessary for the formation of the respiratory supercomplexes.
- Cu stimulates glutathione biosynthesis and glutathione derived specifically from glucose is necessary for survival of Cu*^Hi^* cells. Biosynthesis of glucose-derived glutathione requires activity of glutamyl pyruvate transaminase 2, entry of glucose-derived pyruvate to mitochondria via alanine, and the glutamate exporter, SLC25A22. Glutathione derived from glucose maintains redox homeostasis in Cu-treated cells, reducing Cu-H_2_O_2_ Fenton-like reaction mediated cell death.
- Progression of human ccRCC is associated with gene expression signature characterized by induction of ETC/OxPhos/GSH/Cu-related genes and decrease in HIF/glycolytic genes in subpopulations of cancer cells. Enhanced, concordant expression of genes related to ETC/OxPhos, GSH, and Cu characterizes metabolically active subpopulations of ccRCC cells in regions adjacent to proliferative subpopulations of ccRCC cells, implicating oxidative metabolism in supporting tumor growth.

## INTRODUCTION

Clear cell renal cell carcinoma (ccRCC) is the most frequent renal cancer. The primary treatment for localized disease is surgical removal of primary tumor, however up to 50% of patients relapse within five years after standard-of-care surgical resection. Pembrolizumab is the main approved adjuvant treatment (Choueiri et al., 2021), but currently, there are no predictive or prognostic biomarkers that are used in clinical practice. Thus, there is an urgent clinical need to understand the molecular mechanisms leading to ccRCC relapse and to identify prognostic and predictive biomarkers to guide the adjuvant treatment.

The genetic hallmark of ccRCC is the loss of chromosome 3p, which hosts four major tumor suppressors, *VHL*, *PBRM1*, *SETD2* and *BAP,* and amplification of chromosome 5q (2013; Mitchell et al., 2018; Turajlic et al., 2018). Inactivation of *VHL* and resulting activation of hypoxia inducible transcription factors (HIFs) lead to a pseudohypoxic metabolic phenotype that includes activation of aerobic glycolysis (Warburg effect), pentose phosphate pathway (PPP), serine/glycine biosynthesis pathway, as well as inhibition of pyruvate entry into the TCA (tricarboxylic acid) cycle and an overall decrease in electron transport chain (ETC) and mitochondrial activity (Jonasch et al., 2021; Linehan et al., 2019). Moreover, ccRCC accumulates lipids and glycogen. During ccRCC progression, metabolic reprogramming includes a decrease in lipid content, activation of glutathione metabolism, appearance of dipeptides, a decrease in aspartate levels (Elias et al., 2021; Hakimi et al., 2016), and activation of the TCA cycle (Bezwada et al., 2023). Notably, our recent multiomics analysis of the effects of tobacco smoking, a risk factor for ccRCC, revealed a metabolic shift away from glycolysis towards oxidative phosphorylation (OxPhos), associated with augmented accumulation of Cu and its allocation to cytochrome c oxidase (CuCOX), implicating Cu in the regulation of renal cancer cell metabolism (Reigle et al., 2021).

Cu is a metal cofactor of several enzymes important for cell survival and proliferation, including two catalytic Cu centers in the CuCOX essential for mitochondrial respiration (Chen et al., 2020). High levels of Cu have been reported in sera and tumor tissues in various cancers, while Cu chelators have been developed for clinical trials (Baldari et al., 2020; Ramchandani et al., 2021; Shimada et al., 2018; Wang et al., 2015). Cu-induced OxPhos promotes tumor growth by supporting ATP production (Ishida et al., 2013; Ramchandani et al., 2021; Wang et al., 2015). Other growth effects of Cu include angiogenesis, proliferation, and metalloallosteric regulation of MEK1/2 and ULK1/2 by labile Cu in BRAFV600E driven tumors (Brady et al., 2014; Tsang et al., 2020). Acute exposures of cells to supraphysiological Cu have toxic effects leading to cell death, including a Cu-triggered mitochondrial cell death, termed cuproptosis (Tsvetkov et al., 2022). However, during prolonged exposures, there is an adaptation and resistance to Cu toxicity leading to cuproplasia, i.e. growth effects promoted by Cu (Ge et al., 2022).

Here, we investigated metabolic responses to Cu accumulation in ccRCC growth and progression. We discovered high levels of Cu and increased allocation of Cu to CuCOX in advanced or relapsing tumors. Growth of xenograft tumors was diminished or prevented by low dietary Cu. Chronic exposure to high concentrations of Cu in culture media (Cu*^Hi^*) stimulated bioenergetic, biosynthetic and redox homeostasis pathways. Cu induced mitochondrial oxygen consumption, formation of respiratory supercomplexes and remodeling of cardiolipins. Cu increased PPP activity and de novo biosynthesis of nucleotides. Most importantly, Cu coordinated glucose oxidation with redox homeostasis. Selective abrogation of glucose-derived glutathione production led to cell death dependent on H_2_O_2_ and Cu. Importantly, single cell and spatial transcriptomics of human tumors demonstrated coordinated induction in gene expression of respiratory complexes subunits, and glutathione- and Cu-related genes during progression of ccRCC in subpopulations of metabolically dynamic clones. Tumor regions harboring these metabolically dynamic cancer cells also contained cancer cells characterized by high proliferative state, providing a mechanism by which metabolically active cancer cells can fuel tumor growth. Our data point to the importance of glucose metabolism for maintenance of cellular redox state necessary for survival of cancer cells and identify metabolic pathways as potential vulnerabilities for the development of therapeutic approaches.

## RESULTS

### Cu stimulates progression and growth of ccRCC

We used size exclusion chromatography coupled to inductively coupled plasma mass spectrometry (SEC-ICP-MS) to analyze total Cu content and Cu speciation in primary ccRCCs obtained from the University of Cincinnati (UC) Tumor Bank. Advanced, stage 3+4 tumors had significantly higher levels of total Cu and Cu allocated to CuCOX as compared to stage 1+2 tumors (Figs. 1A and 1B), but no difference in Cu bound to metallothioneins (MTs) or to small molecules in the low molecular weight fraction (LMW) (Figs. 1C and 1D). The content of Cu in CuCOX was significantly correlated with the total Cu concentration only in advanced tumors (Figs. 1E and S1A), suggesting a shift in Cu handling machinery in advanced tumors that favors its allocation to CuCOX over other Cu binding molecules. In contrast, the levels of Cu bound to MTs were significantly correlated with the total Cu concentration independently of the tumor stage, consistent with the role of MTs sequestering a fraction of the total Cu pool. Patients with more advanced tumors also had higher levels of Cu in their sera (Fig. 1F) and there was significant positive correlation in Cu levels between tumor tissues and sera from patients with advanced tumors but not patients with stage 1+2 ccRCCs (Fig. 1G). Note that sera Cu levels in ccRCC patients were higher than the 10-22 µM concentrations observed in unaffected individuals (https://www.ebmconsult.com/articles/lab-test-copper-blood-level), independently of tumor stage.

**Figure 1.**
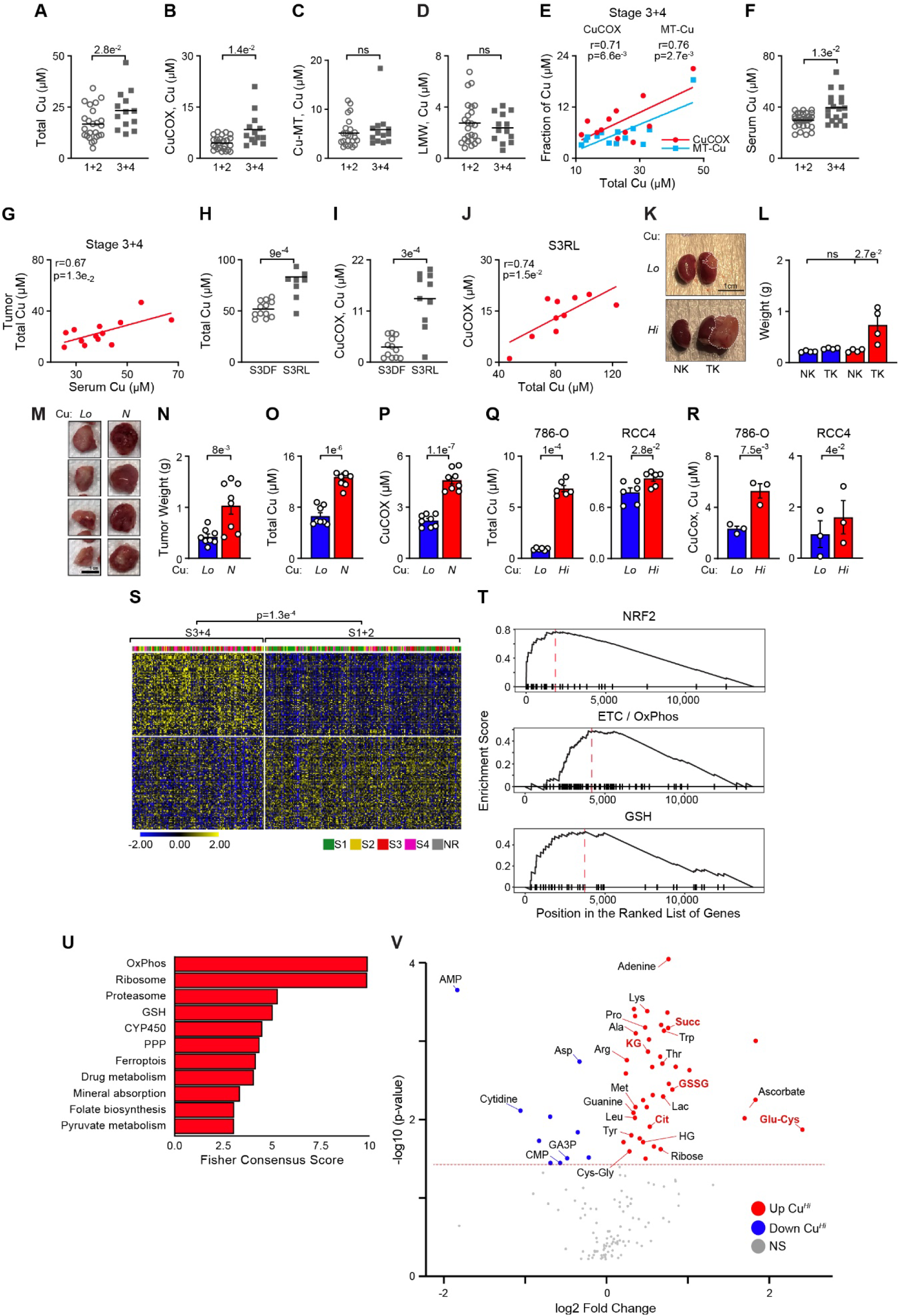
Accumulation of Cu and its allocation to CuCOX associates with progression of ccRCC and drives tumor growth. (A) Total Cu levels in stage 1+2 (n=12) vs. stage 3 (n=26) ccRCCs measured by ICP-MS. (B) Allocation of Cu to cytochrome c oxidase (CuCOX) in stage 1+2 vs. 3+4 tumors (as in A) measured by SEC-ICP-MS. (C) Allocation of Cu to metallothioneins (MTs) in stage 1+2 vs. 3+4 tumors measured by SEC-ICP-MS. (D) Allocation of Cu to low molecular weight fraction (LMW) in stage 1+2 vs. 3+4 tumors measured by SEC-ICP-MS. (E) Pearson correlation of CuCOX and Cu-MT with total tumor Cu levels in 3+4 tumors. (F) Total Cu levels in sera from patients with stage 1+2 tumors vs. 3+4 tumors. (G) Pearson correlation between tumor and serum total Cu levels in stage 3+4 patients. (H) Total Cu levels in tumors from patients with stage 3 ccRCC who relapsed (S3RL, n=10) as compared to those who remained disease free after two years (S3DF, n=12). (I) Cu allocated to CuCOX in tumors from S3RL vs. S3DF patients measured by SEC-ICP-MS. (J) Pearson correlation between CuCOX and total Cu in S3RL tumors. (K) Representative photographs of kidneys with orthotopic xenografts tumors from mice fed matched low (4 µM) and high Cu (158 µM) diet (right) as compared to normal kidneys (left). NK - normal kidney, TK-tumor kidney. Scale bar =1 cm. Tumor tissue is marked by dashed white line. (L) Weight of kidneys with orthotopic xenografts at the time of collection (7 weeks after injection). Cu*^Lo^* - blue, Cu*^Hi^* - red. (M) Representative photographs of subcutaneous xenografts from mice fed standard chow (69 µM, N) and low Cu diet (4 µM, Lo). Scale bar =1 cm. (N) Weight of subcutaneous xenografts at the time of collection (8 weeks after injection). (O) Total Cu levels in xenograft tumors measured by ICP-MS. (P) Allocation of Cu to CuCOX in xenograft tumors measured by SEC-ICP-MS. (Q) Accumulation of total Cu in the indicated cell lines chronically exposed to high Cu in the media. (R) Allocation of Cu to CuCOX measured in the cell lines chronically exposed to high Cu in the media. For RCC4 cells P values calculated by paired t-test. (S) Heatmap shows stratification of ccRCCs from TCGA cohort into clusters enriched in stage 1+2 vs. 3+4 tumors using 200 DEG between Cu*^Hi^* and Cu*^Lo^* 786-O cells. K-means clustering with k=2 was used to cluster both samples and genes. P value from chi square test. (T) Pathways enriched in Cu*^Hi^* cells identified by GSEA. (U) Consensus pathways enriched in both 786-O Cu*^Hi^* cells and metabolically dynamic cancer cell subpopulation 6 identified in scRNA-seq (see also Fig. 6). Fisher consensus of adjusted p-values obtained from individual enrichment analyses is used to rank the pathways. (V) Volcano plot of metabolites’ global levels differentially abundant in Cu*^Hi^* and Cu*^Lo^* 786-O cells. Means ±SEM shown; Unless otherwise indicated, P values from two tailed t-test. Scale bars =1 cm. See also Figure S1, Table S1 and Table S2A.

To test the potential prognostic value of Cu levels in ccRCC, we studied an independent cohort of stage 3 ccRCCs with known disease progression within two years after surgical removal of primary tumors obtained from the UC Tumor Bank. In this independent data set, we identified higher content of total Cu and CuCOX in tumors from patients who relapsed within two years (S3RL) as compared to tumors from patients who remained disease free (S3DF) (Figs. 1H, 1I and S1B), even if the levels of Cu in the sera were not different (Fig. S1C). Importantly, levels of Cu in COX were significantly correlated with total Cu levels in S3RL but not S3DF group (Fig. 1J and S1D). These data support prognostic value of the tumor Cu content.

Next, we sought to determine if dietary exposure to Cu regulates growth of xenografts formed by 786-O RCC cells. Mice were fed custom-made chow based on identical formula except for high and low Cu, or standard chow and custom-made low Cu diet. Low Cu diet significantly inhibited growth of orthotopic (Figs. 1K, 1L, and S1E) and subcutaneous (Figs. 1M, 1N and S1F-H) xenografts. Because of small size, insufficient tumor material could be recovered from orthotopic xenografts grown in mice fed low Cu diet for metallomic analysis. Therefore, we measured Cu and CuCOX in subcutaneous xenografts. Levels of total Cu and Cu allocated to CuCOX were significantly decreased in tumors from mice fed low Cu diet (Figs. 1O, 1P and S1I, S1J). These data point to the activation of a molecular program leading to higher uptake of Cu and its preferential allocation to CuCOX during the progression of ccRCC.

To investigate mechanistic role of Cu in ccRCC metabolic reprogramming, we adapted 786-O and RCC4 renal cancer cells to Cu*^Hi^*conditions. Cells were adapted to high concentration of Cu by gradually increasing Cu concentration in the media over two weeks (until intracellular Cu content reached steady-state) (Fig. S1K). Cu was delivered in a biologically relevant manner, i.e. bound to serum proteins, as verified by SEC-ICP-MS analysis, by preincubation of CuSO_4_ with serum. This allows to deliver Cu in Isogenic Cu*^Lo^*cells were treated identically, except for Cu addition, and cells were continuously cultured in Cu*^Hi^* or Cu*^Lo^* media. The total Cu content (Fig. 1Q) and Cu in CuCOX measured by SEC-ICP-MS (Fig. 1R) were significantly increased in both Cu*^Hi^* cell lines. These cell lines are derived from tumors from white males, a population with the highest incidence of ccRCC (Batai et al., 2019; Feng et al., 2019; Olshan et al., 2013). 786-O cells were able to accumulate more Cu than RCC4 cells, a difference we attribute to the fact that 786-O cells represent more advanced ccRCC, characterized by expression of only HIF2A, while HIF1A is genetically lost (Shen et al., 2011). In contrast, RCC4 cells express both HIFAs. Loss of HIF1A is an event associated with advancement of ccRCC (Hoefflin et al., 2020; Shen et al., 2011).

RNA-seq identified 532 differentially expressed genes (DEG) between Cu*^Hi^* and Cu*^Lo^* 786-O cells (Table S1A). A signature comprising the top 200 differentially expressed genes with the strongest p values stratified the ccRCCs from white males available in the TCGA_KIRC_RNASeqV2_2019 data set (*ilincs*) into two main clusters that were strongly associated with stage (Fig. 1S), supporting role of Cu in tumor progression. Pathway enrichment analysis showed activation of several metabolic pathways, including NRF2, OxPhos, PPP, and glutathione metabolism (Figs. 1T and S1L, Table S1B). Notably, metabolic pathways enrichment in 786-O Cu*^Hi^* cell line showed significant overlap with pathways enriched in a subpopulation of cancer cells with activated oxidative metabolism that we identified in single cell RNA-seq as characterized by concordant activation of glycolysis, ETC and OxPhos activity (Fig. 1U, see also Fig. 6). Consistent with the transcriptomic signature, metabolomic analysis of Cu*^Hi^* and Cu*^Lo^* 786-O cells revealed increased abundance of several intermediates from the TCA cycle and metabolites related to glutathione (Fig. 1V and Table S2A). This metabolic signature is consistent with metabolic profiles identified in advanced ccRCC patient samples (Elias et al., 2021; Hakimi et al., 2016). Overall, these data validate use of RCC cell lines, in particular 786-O Cu*^Hi^* cells, as a suitable model to study biochemical and metabolic effects of Cu in ccRCC.

### Copper remodels ETC activity and mitochondrial biosynthetic pathways

The Cu*^Hi^* cells have augmented mitochondrial oxygen consumption rate (OCR), that includes both basal and maximal respiration, measured by SeaHorse XFe96, and increased ATP production (Fig. 2A-D). This induction of OCR was accompanied by significant biochemical remodeling of ETC respiratory complexes at the level of proteome and lipidome. Using Blue Native-PAGE (Fernandez-Vizarra and Zeviani, 2021), we found that Cu increased formation of the respiratory super complex (RSC) (Fig. 2E) and increased CuCOX activity in this complex measured by in-gel-activity assay (Fig. 2F and 2G) without changes in total protein expression for the representative subunits for each complex (Fig. S2A). Cu induced association of COX17, mitochondrial chaperone delivering Cu to CuCOX (Chen et al., 2020), with lower migrating respiratory complexes (Fig. 2H), supporting preferential allocation of Cu to CuCOX. Moreover, Cu increased expression of CuCOX subunits, (MT-CO1, MT-CO2) and COX7A2L (SCAF1) (Fig. 2I) and there was increased association of COX7A2L with RSC in Cu*^Hi^* cells (Fig. 2J and 2K). This is mechanistically important, as COX7A2L governs formation of RSCs, with functional consequences in response to environmental changes (Fernández-Vizarra et al., 2022; Lapuente-Brun et al., 2013; Pérez-Pérez et al., 2016). In accord with the remodeling of RSCs, LC-MS lipidomics analysis of mitochondrial fractions from Cu*^Hi^* and Cu*^Lo^* 786-O and RCC4 cells revealed a significant increase in cellular cardiolipins (CL), key constituents of mitochondrial membranes associated with formation of RSC (Fig. 2L, S2B, S2C, Table S3). Cardiolipins also were increased in xenograft tumors grown in mice fed high Cu diet as compared to low Cu diet (Fig. 2M and S2D). Chronic Cu exposure had no effect on the overall mitochondrial mass measured by MitoTracker Green in 786-O cells and a very minor increase was measured in RCC4 cells (Fig. S2E). Expression of the mitochondrial protein TOM20 was unchanged by Cu (Fig. S2A).

**Figure 2.**
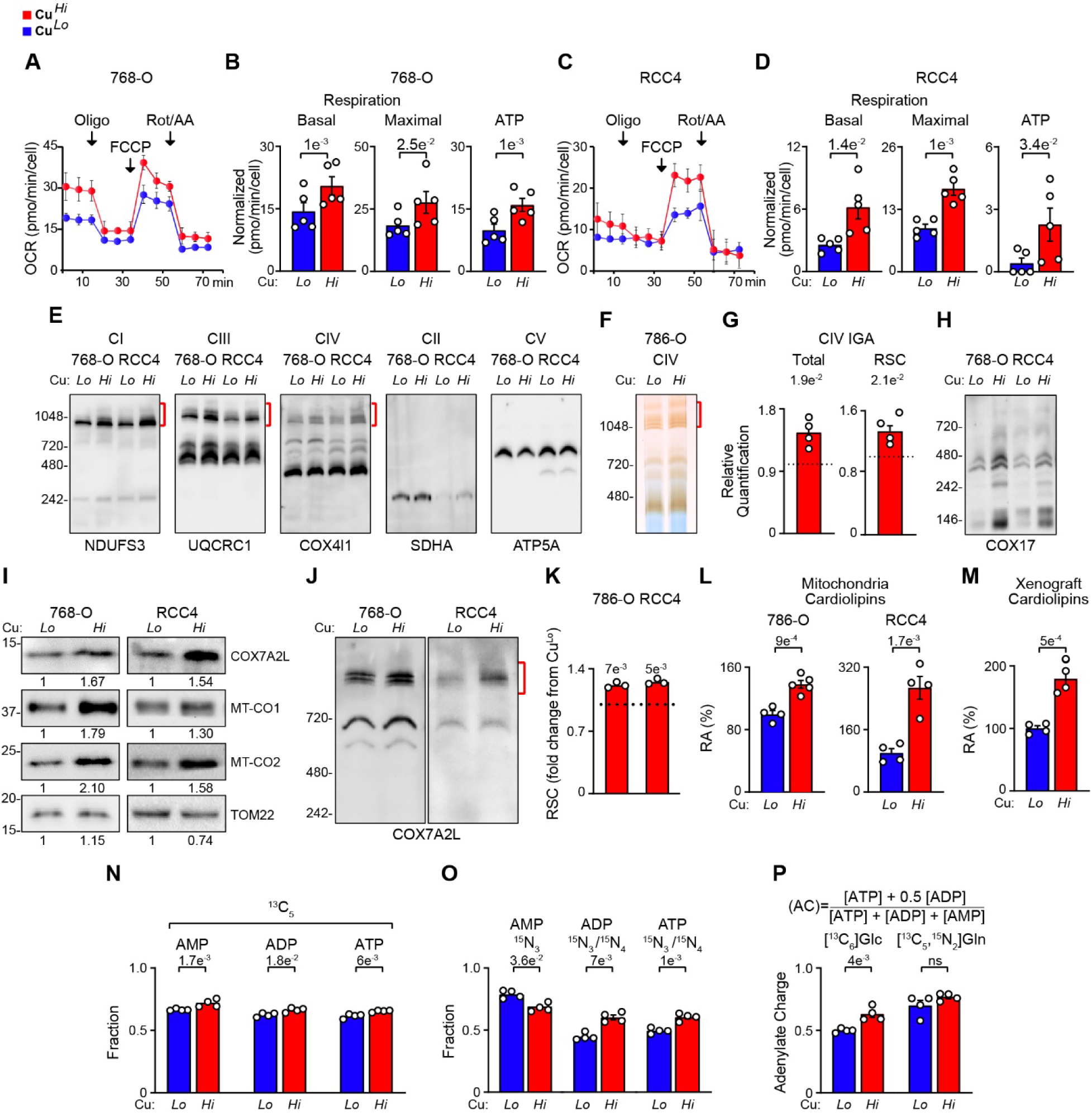
Cu induces mitochondrial oxygen consumption and formation of respiratory supercomplexes. (A-D) Oxygen consumption rate (OCR) from representative seahorse mitochondrial stress tests in 786-O (A, B) and RCC4 (C, D) cells. Quantification of basal respiration, maximal respiration (post FCCP injection), and respiration coupled to ATP production. P value from paired t-test. (E) Blue Native PAGE (BN-PAGE) analysis of respiratory complexes using digitonin permeabilized mitochondria isolated by anti-TOM22 immunopurification from Cu*^Hi^* vs. Cu*^Lo^* cells. Immunoblotting for indicated respiratory complex subunits. Respiratory supercomplex (RSC) is indicated by red brackets. (F) In-gel activity assay (IGA) for cytochrome c oxidase. RSCs indicated with red backet. (G) Quantification of Cu*^Hi^*complex IV (COX) IGA activity relative to Cu*^Lo^*. P values calculated by one-sample t-test. (H) BN-PAGE of mitochondria shows enrichment for COX17, chaperone of Cu to CuCOX. (I) Western blot for COX subunits MT-CO1 and MT-CO2 and RSC assembly factor COX7A2L in mitochondrial lysates. (J) BN-PAGE of mitochondria shows enrichment for COX7A2L in RSCs (red bracket). (K) Quantification of COX7A2L in western blots of RSCs shown in panel M. P values calculated by one-sample t-test. (L) Total cardiolipin content measured by mitochondrial lipidomics in 786-O and RCC4 cells. RA-relative abundance. (M) Total cardiolipin content in xenografts formed by 786-O cells in mice fed low and high Cu diet. RA-relative abundance. (N) Fractions of [^13^C_5_]-adenylate nucleotide labeled from [^13^C_6_]-glucose (Glc) in Cu*^Lo^* and Cu*^Hi^* cells after 24 h of incubation. (O) Fraction of ADP and ATP ^15^N_3_ and ^15^N_4_ labeled form [^13^C_5_,^15^N_2_]-glutamine (Gln) in Cu*^Lo^* and Cu*^Hi^* cells after 24 h incubation. (P) Effect of Cu on adenylate energy charge (AC), formula showed at the top, derived from [^13^C_6_]Glc or from [^13^C_5_,^15^N_2_]Gln in Cu*^Lo^* and Cu*^Hi^*cells after 24 h of incubation. Means ±SEM shown; unless indicated, P values were calculated from unpaired two tailed t-test. See also Figure S2 and Table S3.

Next, we analyzed effects of Cu on the contribution of glucose and glutamine, major carbon and nitrogen donors, to nucleotides using flux of ^13^C and ^15^N from [^13^C_6_]-glucose or [^13^C_5_,^15^N_2_]-glutamine (Table S2B and S2C). Cu significantly increased the fraction of the ^13^C_5_ labeled adenylate nucleotides in the total pool of each nucleotide, an indication of increased utilization of ribose derived from glucose via the PPP (Fig. 2N) and increased flux of ^13^C into 6-phospho-D-gluconate indicating activation of PPP (Fig. S2F). This is consistent with transcriptomic data demonstrating increased PPP gene expression in Cu*^Hi^* cells (see Fig. S1L and 1U). Cu increased the fractions of labeled ADP and ATP (Fig. 2O) and labeled GTP, UTP and CTP (Fig. S2G) in the total pool of each nucleotide after 24 h incubation with labeled glutamine. These data show effects of Cu on major anabolic pathways supporting its overall oncogenic activity.

Cellular energy status that represents metabolic requirements is best represented by adenylate energy charge (AC), which considers the concentrations of all three adenine nucleotides, ATP, ADP and AMP (Fig. 2P) (Atkinson and Walton, 1967; Swedes et al., 1975). Interestingly, Cu augmented AC derived from [^13^C_6_]-glucose when calculated using [^13^C_5_]ATP, ADP, and AMP (Fig. 2P). In contrast, Cu had no effect on AC when calculated using ^15^N_3_ and ^15^N_4_ in adenylate nucleotides derived from [^15^N_2_]-glutamine (Fig. 2P). This suggests that Cu differently regulates contributions of glucose and glutamine to energy homeostasis.

Overall, these data indicate that Cu stimulates bioenergy production by activating mitochondrial utilization of glucose in a dual mechanism that includes its oxidation and biosynthesis of adenylate nucleotides, accompanied by comprehensive protein and lipid remodeling of ETC complexes.

### Copper induces dependence on glucose-derived glutathione for survival

To further understand the functional role of glucose or glutamine metabolism in Cu*^Hi^* and Cu*^Lo^* cells, we determined the effects of glucose or glutamine starvation on cell survival. Cu*^Hi^* cells were markedly more sensitive to glucose starvation than Cu*^Lo^* cells (Fig. 3A), an effect largely rescued by exogenous pyruvate or dimethylketoglutarate (DMKG) (Fig. 3B). In contrast, glutamine starvation induced a similar degree of cell death in both Cu*^Hi^* and Cu*^Lo^*cells (Fig. S3A), partially rescued by DMKG but not pyruvate (Fig. S3B). In parallel studies, we observed that Cu*^Hi^* cells were also selectively hypersensitive to the inhibition of glutathione biosynthesis with buthionine sulphoximine (BSO) (Nishizawa et al., 2018) (Fig. 3C). Moreover, GSH and GSSG levels were higher in Cu*^Hi^* as compared to Cu*^Lo^* cells (Fig. 3D), complementing and functionally validating metabolomic data showing increased abundance of intermediates of glutathione pathway in Cu*^Hi^* cells (Fig. 1V, Table S2A). Next, we examined the effects of glucose and glutamine starvation on cellular glutathione levels. Glucose starvation decreased only GSH but not GSSG in Cu*^Hi^* but not in Cu*^Lo^*cells (Fig. 3D). GSH/GSSG ratio was lower in Cu*^Hi^* cells as compared to Cu*^Lo^*cells under normal glucose condition, and it was further decreased by glucose starvation only in Cu*^Hi^* cells. (Fig. S3C). In contrast, glutamine starvation led to substantial and similar depletion of both GSH and GSSG in Cu*^Lo^* and Cu*^Hi^* cells (Fig. 3D). The critical role of glucose-derived glutathione in survival of Cu*^Hi^* cells was supported by the fact that exogenously supplied ethyl ester of GSH (GSH-EE) significantly prevented cytotoxicity induced by glucose starvation (Fig. 3E), while BSO prevented rescue of the cytotoxic effect of glucose starvation by exogenous pyruvate (Fig. 3F). Moreover, exogenous pyruvate recovered the GSH/GSSG ratios in glucose starved Cu*^Hi^* cells (Fig. 3G). In contrast, cell death induced by glutamine starvation in Cu*^Hi^* and Cu*^Lo^* cells was not prevented by exogenous GSH-EE (Fig. S3B).

**Figure 3.**
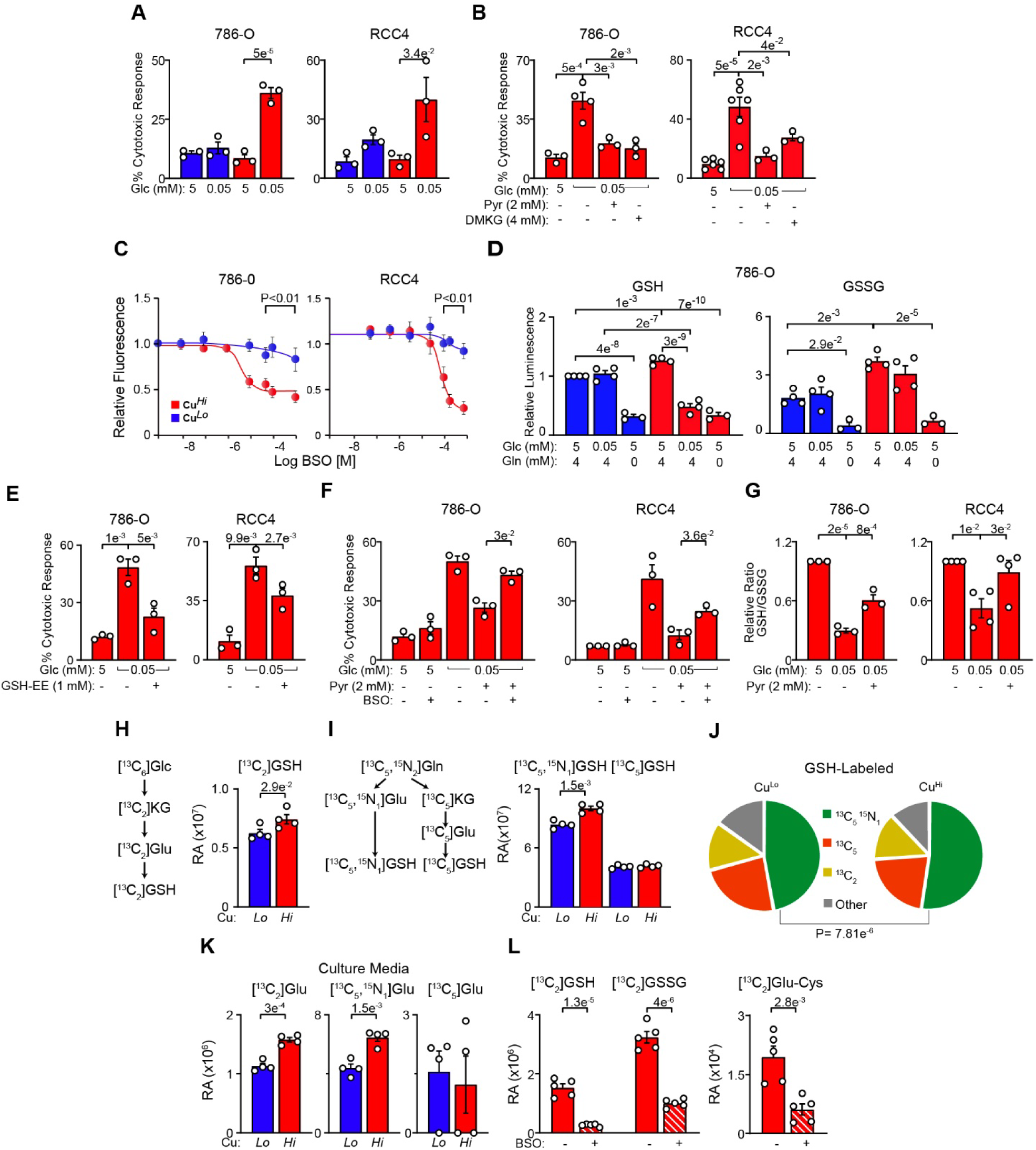
Glucose-derived glutathione promotes survival of Cu*^Hi^* cells. (A) Cytotoxic effect of glucose (Glc) starvation on 786-O and RCC4 Cu*^Hi^* but not Cu*^Lo^* cells measured using propidium iodide and FITC Annexin V staining and flow cytometry. (B) Rescue of glucose starvation induced Cu*^Hi^* cell death with exogenous pyruvate (Pyr) and dimethylketoglutarate (DMKG) in 786-O and RCC4 cells. (C) Cu*^Hi^* but not Cu*^Lo^* cells are sensitive to inhibition of glutathione biosynthesis by BSO (48 h) measured by CyQuant proliferation assay. Because EC_50_ for BSO dose in Cu*^Lo^* cells could not be determined, the P values were determined for each set of data points in each cell line using two-tailed t-test. (D) Effects of glucose and glutamine starvation on GSH and GSSG levels in Cu*^Hi^* and Cu*^Lo^*cells measured using GSH-Glo Assay. (E) Rescue of glucose starvation-induced Cu*^Hi^* cell death with exogenous GSH ethyl ester (GSH-EE). (F) Inhibition of glutathione biosynthesis with BSO (16 h) reverses pyruvate induced rescue of Cu*^Hi^* cells death caused by glucose starvation. P values calculated from two-tailed t-test. BSO concentrations: 10 μM (786-O) and 25 μM (RCC4). (G) Exogenous pyruvate reverses glucose starvation induced decrease in GSH/GSSG ratio. (H) Schematic pathway of ^13^C labeling of GSH derived from [^13^C_6_]Glc and relative abundance (RA) in [^13^C_2_]GSH in 786-O cells after 24 h incubation with labeled glucose. P value from two tailed t-test. (I) Schematic pathway of ^13^C and ^15^N labeling of GSH derived from [^13^C_5_,^15^N_2_]Gln and relative abundance (RA) of [^13^C_5_,^15^N_1_]GSH and [^13^C_5_]GSH pools in 786-O cells after 24 h incubation with labeled glutamine. P values from two tailed t-test. (J) Fractions of [^13^C_5_,^15^N_1_]GSH, [^13^C_5_]GSH and [^13^C_2_]GSH, in Cu*^Lo^* and Cu*^Hi^* 786-O cells. P value from two tailed t-test. (K) Increased levels of [^13^C_2_]Glu and [^13^C_5_,^15^N_1_]Glu but not [^13^C_5_]Glu in the culture media of Cu*^Hi^* 786-O cells after 24 h of incubation with labeled metabolite. RA-relative abundance. P values from two tailed t-test. (L) BSO inhibits ^13^C_2_ labeling of glutathione metabolites from [^13^C_2,3_]Pyr. RA-relative abundance. P values from two tailed t-test. Means ±SEM shown; unless indicated, P values are from one-way ANOVA with Holm-Sidak post-test. See also Figure S3 and Table S2.

Using metabolic labeling with [^13^C_6_]-glucose or [^13^C_5_,^15^N_2_]-glutamine we identified three sources of labeled intracellular glutathione. [^13^C_2_]GSH was the predominant glucose-derived form of labeled GSH in cells treated with labeled glucose and its abundance was increased in Cu*^Hi^* cells contributing 14% of the total GSH in Cu*^Hi^* and Cu*^Lo^* cells (Fig. 3H and 3J). Both the abundance and the fraction of glutamine derived [^13^C_5_,^15^N_1_]GSH were increased in Cu*^Hi^* cells, and represented the majority of the total GSH (Fig. 3I and 3J). In contrast, the abundance and fraction of glutamine-derived [^13^C_5_]GSH were not changed by Cu and contributed about 20% of total cellular GSH (Fig. 3I and 3J). In accordance with GSH labeling, chronic Cu treatment increased excretion into the culture media of glucose-derived [^13^C_2_]Glu and glutamine-derived [^13^C_5_,^15^N_1_]Glu, but not [^13^C_5_]Glu (Fig. 3K). This indicates that utilization of the Glu derived from different sources for Cu-induced GSH biosynthesis is matched by Glu utilization for cystine exchange at the plasma membrane. Finally, we validated the flux of carbon from pyruvate to GSH by measuring enrichment of intracellular GSH, GSSG and Ɣ-glutamyl-cysteine, when glucose starved cells were treated with [2,3-^13^C_2_] pyruvate, an effect prevented by co-treatment with BSO (Fig. 3L). Importantly, fractions of these [^13^C_2_] labeled glutathione metabolites derived from ^13^C labeled pyruvate were similar to the fraction of ^13^C_6_ glucose derived [^13^C_2_]GSH after 24 h of labeling and they were respectively at 11.6%, 12.1% and 16.1%. We also detected 14% of [^13^C_2_]KG and 17% of [^13^C_2_]Glu from labeled pyruvate, supporting the role of the TCA cycle in biosynthesis of a pool of glutathione from glucose.

Glutathione biosynthesis is regulated by the NRF2 transcription factor (He et al., 2020) and Cu*^Hi^*cells showed induction of NRF2 signature (Fig. 1T). Cu*^Hi^* cells exhibited increased nuclear localization of NRF2 accompanied by decreased cytosolic expression of its inhibitory partner, KEAP1 (Fig. S3D), as well as increased expression of NRF2 target genes including catalytic and modulatory subunits of glutamate-cysteine ligase (GCLC and GCLM), the rate limiting enzyme in glutathione biosynthesis, and SLC7A11, an amino acid antiporter that exchanges extracellular cystine for intracellular glutamate (Fig. S3E and S3F).

These results show biochemical compartmentalization and different functional contribution of glucose- and glutamine-derived glutathione and demonstrate the selective recruitment of a relatively small fraction of glucose-derived reduced glutathione to maintain viability of Cu*^Hi^* cells.

### Glutamate pyruvate transaminase 2 and the mitochondrial glutamate carrier SLC25A22 are required for biosynthesis of glucose-derived glutathione and survival of Cu*^Hi^* cells

Glucose-derived pyruvate is used in several biochemical reactions. In the cytosol, it is converted to lactate by lactate dehydrogenase (LDH), regenerating the NAD needed for active glycolysis. It enters the mitochondria via the pyruvate carriers (MPC1/MPC2) to be converted to acetyl-CoA by pyruvate dehydrogenase (PDH) complex. Pyruvate is also converted to alanine by glutamate pyruvate transaminase (GPT) (Fig. 4A). In ccRCC, conversion of pyruvate to acetyl-CoA is inhibited by HIF-induced PDH kinase (PDK1) that phosphorylates PDHA subunit and inhibits PDH activity (Kim et al., 2006; Papandreou et al., 2006).

**Figure 4.**
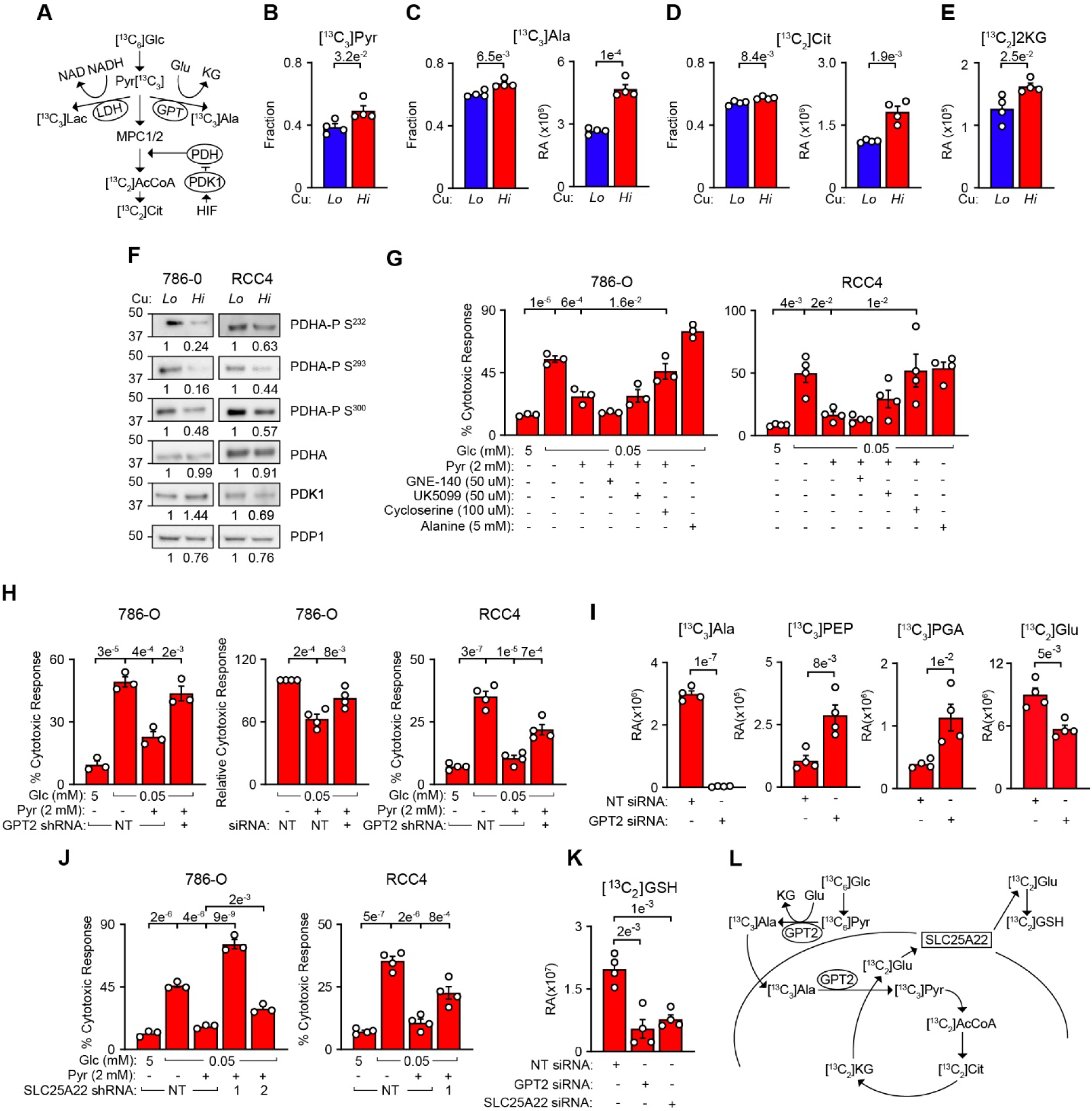
Synthesis of glucose derived glutathione requires activity of glutamyl pyruvate transaminase 2 and mitochondrial glutamate carrier SLC25A22. (A) Pathways of pyruvate utilization (B) Fraction of [^13^C_3_]Pyr in Cu*^Lo^* and Cu*^Hi^* 786-O cells treated with [^13^C_6_]Glc for 5 h. (C) Fraction and relative abundance (RA) of [^13^C_3_]-alanine (Ala) in Cu*^Lo^* and Cu*^Hi^* 786-O cells treated with [^13^C_6_]Glc for 24 h. (D) Fraction of total and relative abundance (RA) of [^13^C_2_]-citrate (Cit) in Cu*^Lo^* and Cu*^Hi^* 786-O cells treated with[^13^C_6_]Glc for 24 h. (E) Relative abundance of [^13^C_2_]-ketoglutarate (KG) in Cu*^Lo^* and Cu*^Hi^* 786-O cells with [^13^C_6_]Glc for 24 h. (F) Western blot showing decrease in phosphorylation of PDHA at the indicated serine residues in Cu*^Hi^* 786-O and RCC4 cells. (G) Effects of pharmacological treatments on pyruvate rescue of cell death induced by glucose starvation in Cu*^Hi^* 786-O and RCC4 cells. GNE-140, LDH inhibitor; UK5099, MPC1/2 inhibitor; Cycloserine, GPT/GPT2 inhibitor. (H) Effects of GPT2 knockdowns on rescue of glucose starvation induced cell death by exogenous pyruvate in 786-O and RCC4 cells. NT-non-targeting sh or siRNA. (I) Effects of GPT2 knockdown on labeling of [^13^C_3_]-Ala, -PEP and -PGA and [^13^C_2_]Glu in 786-O cells treated with [^13^C_6_]Glc for 24 h. P values from two tailed t-test. (J) Effects of SLC25A22 knockdown on pyruvate dependent rescue of glucose starvation induced cell death in 786-O and RCC4 cells. Data shown in the first three bars for RCC4 cells are the same as in panel H. (K) Effects of GPT2/SLC25A22 knockdownss on flux of carbon from [^13^C_6_]Glc to [^13^C_2_]GSH. P value from two-tailed t-test. (L) Model of the proposed pathway. In all figures Mean±SEM is shown. In all n≥ 3. P values from two-tailed t-test in B-E, I or one-way ANOVA with Holm-Sidak post-test in G, H, J. See also Figure S4 and Table S2B.

Stable isotope tracing using [^13^C_6_]-glucose in Cu*^Hi^* cells revealed a significantly increased fraction of [^13^C_3_]-pyruvate after 5 h of labeling (Fig. 4B), and increased flux of glucose carbon into all three products downstream from pyruvate. There was a significant increase in labeled [^13^C_3_]-lactate after 5 h (Fig. S4A) and an increased abundance and fraction of [^13^C_3_]-alanine after 24 h of incubation with labeled glucose (Fig. 4C). Moreover, after 24 h, there was a significant increase in the abundance and fraction of [^13^C_2_]-citrate (Fig. 4D) and abundance of [^13^C_2_]-ketoglutarate (Fig. 4E), indicating an increased entry and processing of glucose carbon via the TCA cycle. These data showing increased flux of pyruvate to the TCA cycle via PDH indicate that the inhibitory effect of PDK1 on PDH activity is decreased in Cu*^Hi^* cells. In line with this, in Cu*^Hi^* cells we found decreased phosphorylation of PDHA Ser232, 293 and 300 without major changes in the expression of PDK1 or phosphatase, PDP1 (Fig. 4F).

Next, we investigated the functional role of pathways utilizing pyruvate in the rescue of glucose starvation dependent cell death. Inhibition of LDH with the selective inhibitor GNE-140 (Laganá et al., 2019) slightly increased pyruvate’s rescue of glucose starvation induced cytotoxicity in 786-O cells but had no effect in RCC4 Cu*^Hi^* cells (Fig. 4G). Inhibition of MPC1/2 pyruvate transporters with UOK5099 did not affect pyruvate rescue during glucose starvation in neither cell line. In many cancers, including ccRCC, the activity of MPC1/2 pyruvate transporters is low and low MPC levels are associated with poor prognosis (Ruiz-Iglesias and Mañes, 2021; Tang et al., 2019) (Fig. S4B). This result points to the alternative pathways of pyruvate entry into mitochondria. Importantly, an inhibitor of glutamate pyruvate transaminases (GPT/GPT2), cycloserine (Cornell et al., 1984) reversed pyruvate rescue of glucose starvation-induced cell death (Fig. 4G), an indication that pyruvate may enter mitochondria via alanine transport. Expression of the predominantly cytosolic GPT is very low in renal cancer cells (The Human Protein Atlas) and was not detected in 786-O and RCC4 cells (Fig. S4C), while mitochondrial GPT2 can also be cytosolic (Fig. S4D). Genetic inhibition of GPT2 using si and validated shRNA (Sousa et al., 2016; Wei et al., 2022) reversed pyruvate rescue of glucose starvation induced cell death (Figs. 4H and S4E). The effectiveness of the GPT2 knockdown was further validated by complete lack of [^13^C_3_]-alanine and increased accumulation of the glycolytic metabolites [^13^C_3_]-phosphoenolpyruvate (PEP) and -phosphoglycerate (PGA), indicating inhibition of the pathway using glucose carbon for alanine biosynthesis (Fig. 4I). GPT2 knockdown significantly decreased the abundance of [^13^C_2_]Glu, an indication that GPT2 was required to generate glucose-derived glutamate (Fig. 4I). Expression levels of GPT2 were minimally affected in Cu*^Hi^* as compared to Cu*^Lo^* cells (Fig. S4C). Importantly, addition of alanine to the culture media did not reverse glucose starvation-induced cell death (Fig. 4G), demonstrating that the activity of GPT2 but not production of alanine was required for the rescue. We propose that in the cytosol, pyruvate undergoes transamination to alanine and alanine enters mitochondria where it is converted back to pyruvate. GPT2 activity is then further required for selective transamination of glucose-derived [^13^C_2_]KG to [^13^C_2_]Glu, which is used in GSH biosynthesis essential for survival of Cu*^Hi^* cells.

Glutathione biosynthesis is localized to the cytosol, thus glucose-derived [^13^C_2_]Glu needs to exit mitochondria. Two mitochondrial glutamate carriers, SLC25A22 and SLC25A18 (GC1 and GC2) show bidirectional activity symporting glutamate and H^+^ to and from mitochondria (Fiermonte et al., 2002; Frigerio et al., 2008; Schoolwerth et al., 1983). We prioritized analysis of SLC25A22 because its high mRNA level is associated with significantly worse survival in ccRCC (Fig. S4F). Knockdown of the SLC25A22 transporter reversed rescue of glucose starvation induced cytotoxicity by exogenously provided pyruvate (Fig. 4J and S4G), indicating that export of glutamate from mitochondria by SLC25A22 is essential for cell survival in Cu*^Hi^* cells. Furthermore, we determined that knockdowns of GPT2 and SLC25A22 significantly diminished flux of carbon from [^13^C_6_]-glucose into [^13^C_2_]GSH (Fig. 4K), supporting the role of this pathway in the biosynthesis of a pool of glutathione essential for survival of Cu*^Hi^* cells.

Overall, these data support the role of GPT2 and SLC25A22 in production and delivery of mitochondrial glutamate derived from glucose to be used in the cytosol for biosynthesis of a pool of glutathione (Fig. 4L). This mechanism couples biosynthesis of glutathione and regulation of redox homeostasis with glucose oxidation and ETC activity, a process that is essential for survival of Cu*^Hi^* cells.

### The glucose-regulated glutathione pool preserves redox homeostasis in Cu*^Hi^* cells

Cu*^Hi^* cells show constitutive oxidative stress as demonstrated by reduced GSH/GSSG ratio (Fig. S3C). Glucose starvation further lowered GSH/GSSG ratios in Cu*^Hi^* cells (Fig. S3C and 3G). Glucose starvation induced OCR in both Cu*^Lo^* and Cu*^Hi^* cells, the effect was stronger in Cu*^Hi^* cells, and was not reversed by exogenous pyruvate treatment (Fig. 5A). The increase in OCR was accompanied by augmented mitochondrial superoxide accumulation measured by MitoSOX that was significantly higher in Cu*^Hi^* cells and was reversed by pyruvate (Fig. 5B). Using the CM-H2DCFDA probe that is more specific for H_2_O_2_, we measured an increase in H_2_O_2_ in response to glucose starvation only in Cu*^Hi^*cells and this effect was completely reversed by pyruvate (Fig. 5C). Furthermore, the pyruvate rescue of H_2_O_2_ accumulation in glucose starved cells was prevented by inhibition of glutathione biosynthesis with BSO (Fig. 5C) or knockdowns of GPT2 (Fig. 5D) and SLC25A22 (Fig. 5E). In line with this, treatment with exogenous GSH-EE inhibited accumulation of H_2_O_2_ (Fig. 5F).

**Figure 5.**
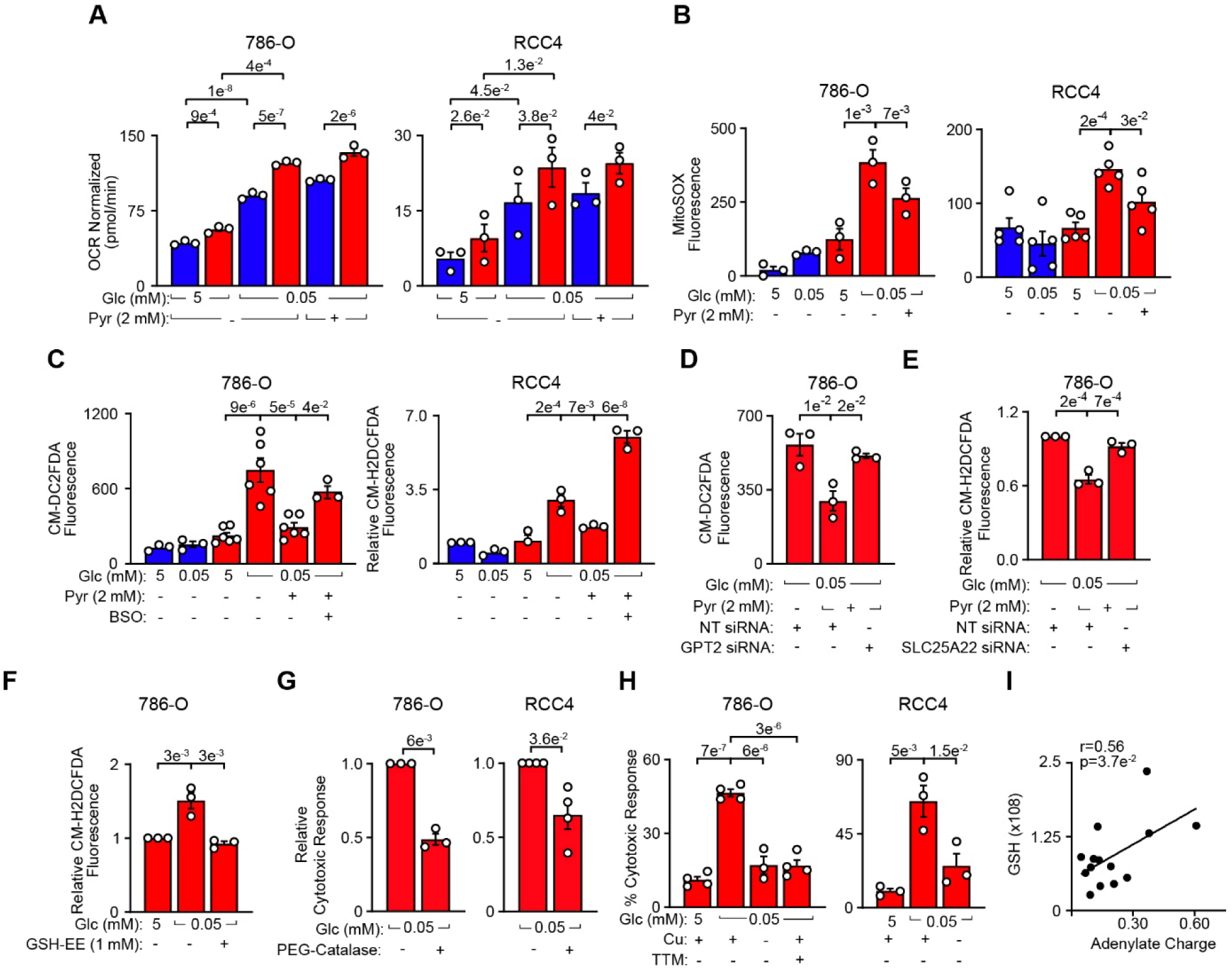
Glucose-derived mitochondrial glutathione prevents Fenton reaction-like mediated cell death of Cu*^Hi^* cells. (A) Seahorse measurement of basal OCR in response to glucose starvation in Cu*^Lo^* and Cu*^Hi^* 786-O and RCC4 cells. Data are normalized to Hoechst staining. (B) MitoSOX measurement of mitochondrial superoxide accumulation in response to glucose starvation (16 h) and treatment with exogenous pyruvate in Cu*^Lo^* and Cu*^Hi^* 786-O (P value from paired t-test) and RCC4 (P values from unpaired t-test) cells. (C) CM-H2DCFDA measurement of cellular hydrogen peroxide in response to glucose starvation and treatments with exogenous pyruvate and BSO in Cu*^Lo^* and Cu*^Hi^* 786-O and RCC4 cells. BSO concentrations: 10 µM for 786-O and 25 µM for RCC4 cells. (D) Effect of GPT2 knockdown on the accumulation of H_2_O_2_ in response to pyruvate rescue of glucose starvation in Cu*^Hi^* cells. (E) Effect of SLC25A22 knockdown on the accumulation of H_2_O_2_ in response to pyruvate rescue of glucose starvation in Cu*^Hi^*cells. (F) Effect of exogenous glutathione GSH-EE on the accumulation of H_2_O_2_ in response to glucose starvation in Cu*^Hi^* cells. (G) Cell death caused by glucose starvation is prevented by treatment with PEG-catalase. 786-O cells: 250 U/ml, RCC4 cells 500 U/ml. P-values from two-tailed t-test. (H) Glucose starvation induced death of Cu*^Hi^*cells prevented by removal of Cu or treatment with Cu chelator (TTM, 30 µM) in 786-O and RCC4 cells. (I) Positive Pearson correlation between GSH relative abundance and adenylate charge measured in a cohort of 16 stage 3 ccRCCs. Means ±SEM shown; unless otherwise indicated, P values calculated from one-way ANOVA with Holm-Sidak post-test. See also Figure S5.

Expression levels of enzymes detoxifying ROS, such as SOD1, SOD2, catalase, mitochondrial GSH-independent peroxiredoxin 3 (PRDX3), cytosolic GSH-dependent peroxiredoxin 6 (PRDX6) were not affected by Cu or glucose concentration, while expression of mitochondrial glutathione peroxidases (GPX1/2) was decreased in Cu*^Hi^* cells, an effect that may be attributed to the known activity of Cu in reducing expression of cellular selenoproteins (Schwarz et al., 2020; Xue et al., 2023) (Fig. S5). These results indicate that GSH is involved in alternative mechanisms preventing accumulation of H_2_O_2_, by modulating H_2_O_2_ generation or decomposition, or both.

Next, we investigated the mechanism of cell death induced by glucose starvation. Treatment of glucose starved Cu*^Hi^*cells with PEG-catalase prevented death, supporting a causative role of H_2_O_2_ in death of Cu*^Hi^* cells (Fig. 5G). Surprisingly, glucose-starvation induced cell death was completely rescued by Cu-free media applied for the duration of starvation or by tetrathiomolybdate (TTM), an intracellular Cu chelator (Lowndes et al., 2009) (Fig. 5H). These data are consistent with Fenton-like chemistry involving the labile Cu and H_2_O_2_ in cell death during glucose deprivation in Cu*^Hi^* cells. Fenton reactions generate hydroxyl radicals that mediate broad oxidative stress causing cell death via oxidation of nucleic acids, lipids and proteins.

Overall, these data indicate that chronic adaptation to high Cu requires contribution of glucose-derived reduced glutathione that is used to limit accumulation of H_2_O_2_ generated by ETC activity to prevent toxicity of Fenton-like reaction with Cu. Absence of this GSH activity results in cell death. This coordinated regulation of energy production and redox homeostasis is further supported by a significant positive correlation between levels of GSH and adenylate charge measured in stage 3 ccRCCs (Fig. 5I).

### Activation of a metabolic state characterized by high ETC/OxPhos, Cu and GSH metabolism is a hallmark of ccRCC progression

To determine the activity of ETC/OxPhos/Cu/GSH pathways during progression of ccRCC, we investigated expression of genes encoding subunits of the ETC, as well as Cu and glutathione related genes in three independent cohorts of tumors using single cell, bulk, and spatial transcriptomics data. Meta-analysis of single cell transcriptomic data of 18 ccRCCs from two established cohorts (Li et al., 2022; Obradovic et al., 2021) was used to identify subpopulations of cancer cells (Fig. 6A). Cancer cells were identified by expression of the HIF target, carbonic anhydrase 9 (CA9), (Fig. S6A) and confirmed by inferred copy variant number analysis showing loss of chromosome 3p and gain of chromosome 5q (Fig. S6B), the two most frequent genomic alterations in ccRCC (Mitchell et al., 2018). The cancer cells were subsequently clustered by their transcriptional similarity, and the stability of cell subpopulations was assessed using repeated re-clustering with randomly selected 80% of cells. The subpopulations, 2, 6 and 10, were identified as most stable with the median of Max Jaccard Index > 0.6 (Tang et al., 2021) (Fig. 6B) and subsequently characterized in terms of their distinct metabolic and proliferative states.

**Figure 6.**
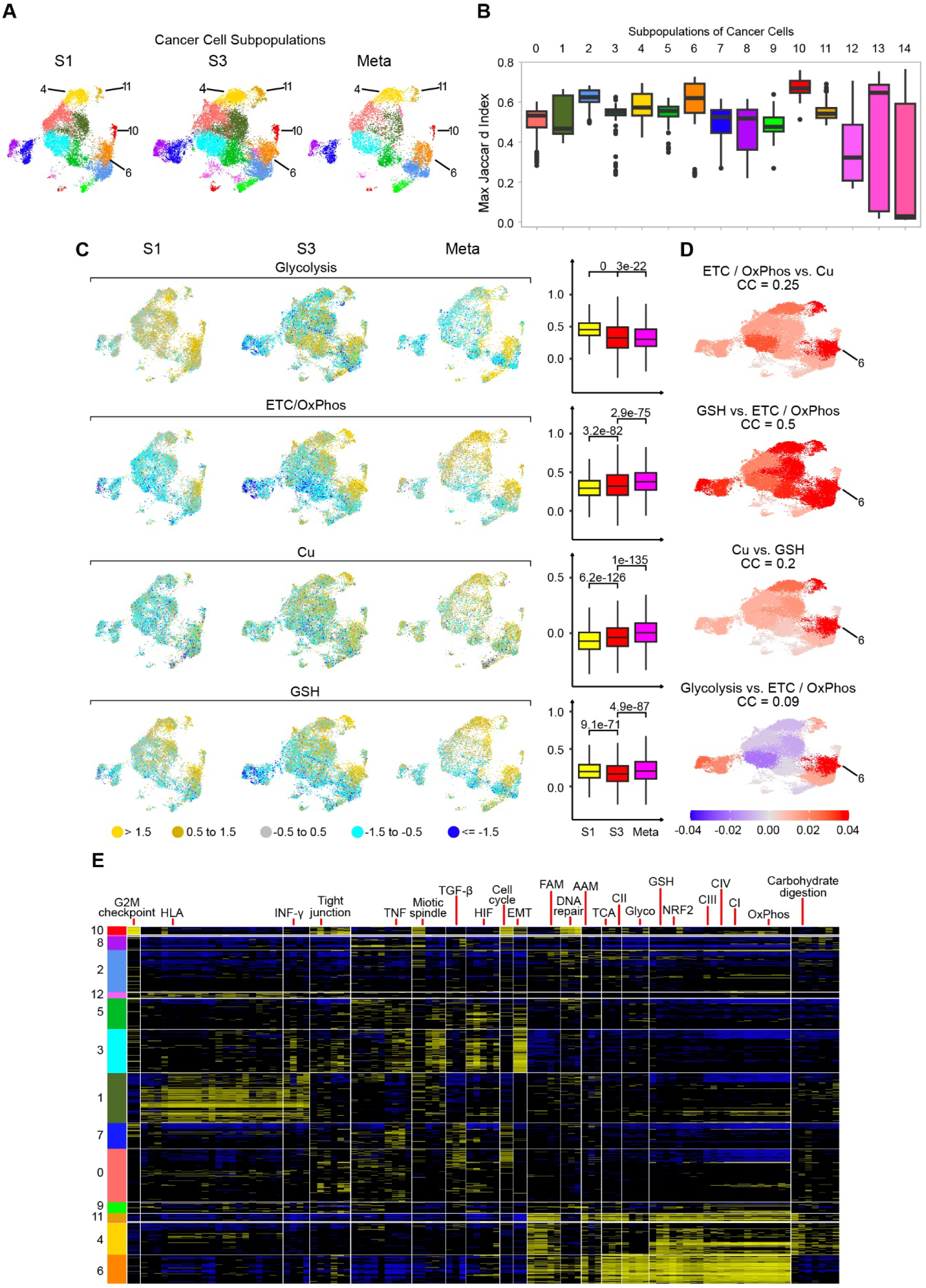
Induction of ETC/OxPhos, Cu and GSH related genes during progression of ccRCC. (A) UMAP visualization of 15 subpopulations of cancer cells. S1 - stage 1 (n=6 tumors and 13,689 cells); S3 - stage 3 (n=9 tumors and 19,273 cells); Meta – metastatic (n=3 tumors and 9,072 cells). (B) Resampling analysis of cluster stability using Jaccard Index (stability score) for each subpopulation of cancer cells, with subpopulations 2, 6 and 10 showing the highest levels of stability. (C) UMAPs (left) show overall decrease in expression of genes encoding glycolytic genes while induction of ETC/OxPhos, Cu and GSH related genes in more advanced ccRCCs. Box-whisker plots (right) show the distributions of module scores for each pathway signature across all putative cancer cells. Each column shows expression of the indicated gene set using the Seurat module score defined relative to a control gene set. (D) Pearson correlation residuals of the indicated sets of genes in each subpopulation of cancer cells shown for all tumors (n=18) combined. (E) Patterns of pathways expression in cancer cell subpopulations. Rows represent cells, with clusters of cells identified using Seurat analysis of scRNA-seq data indicated by vertical color bars and sorted according to their overall similarity using cluster centroids. Columns represent supergenes and pathways, selected from the union of KEGG pathways, MSigDB Hallmark gene sets, and curated metabolic gene sets. Supergenes/pathways are clustered based on their similarity using 1-Pearson correlation coefficient as the dissimilarity measure. Expression values for a supergene/pathway in a cell are defined as the average normalized expression over all genes in the set and shown using 3 state projection: top quartile (high expression) in yellow, bottom quartile (low expression) in blue, middle 50% (average expression) in black. EMT – epithelial mesenchymal transition; FAM – fatty acid metabolism; AAM – amino acid metabolism; CI/CII/CIII/CIV - respiratory complexes I/II/III/IV; Glyco – glycolysis. See also Figure S6 and Table S4.

Overall, there was a clear, stage-dependent global transcriptional reprogramming indicating reduction of a glycolytic phenotype and enhancement of oxidative metabolism during progression of ccRCC. Expression of ccRCC’s glycolytic genes and HIF targets was decreased in stage 3 and metastatic tumors as compared to stage 1 tumors in several subpopulations of cancer cells (Figs. 6C and S6C). In contrast, the expression of a superset of genes encoding all subunits of mitochondrial respiratory complexes (ETC/OxPhos) or individual complexes, genes associated with Cu allocation and genes related to glutathione metabolism were increased in more advanced tumors (Fig. 6C and S6C, Table S4). Importantly, there was induction of a NRF2 transcriptomic signature (Fig. S6C), which is in line with the established role of this transcription factor in regulation of glutathione biosynthesis and ROS detoxification (DeNicola et al., 2011; He et al., 2020) and with our cell line data (Figs. 1T and S3D-F). The expression levels of gene sets in metabolic pathways for ETC/OxPhos, Cu and GSH were significantly correlated across all cell subpopulations (Fig. 6D). Interestingly, there was lack of concordance in expression of ETC/OxPhos and glycolytic genes in the majority of subpopulations, except for subpopulation 6, where both sets of genes were expressed together (Fig. 6D). Visualization of expression patterns for metabolic and other gene sets further revealed strong concordant expression of glycolytic, ETC, glutathione and NRF2 related genes in subpopulation 6 (Fig. 6E). This implicates activation of full glucose oxidation accompanied by induction of glutathione redox scavenging system. Subpopulations 4 and 11 were also characterized by induction of metabolic genes, including ETC, OxPhos, TCA cycle, fatty acid metabolism, but not glycolysis, implicating non-glucose derived entry of electrons into ETC (Fig. 6E). Note that subpopulation 6 was transcriptionally similar to the cell culture model, 786-O Cu*^Hi^* cells (Fig. 1U). Stable subpopulation 10 was noteworthy for higher expression of cell cycle and DNA repair genes indicating proliferative phenotype (Fig. 6E). This subpopulation showed also increased expression of OxPhos related genes (Fig. 6E).

The significance of the identified mitochondrial phenotype in progression of ccRCC was further corroborated by the bulk transcriptomic analysis of stage 3 ccRCCs from the TCGA KIRC Firehose Legacy cohort. We focused on patients with known information about progression, i.e., patients who either remained disease free (S3DF) or relapsed (S3RL) within two years from the resection of the primary tumor (Fig. S7A). Note that tumors from S3RL patients showed significantly higher levels of total Cu and CuCOX as compared to tumors from S3DF patients (Figs. 1H-1J). We identified 1,267 differentially expressed genes of which 340 and 927 were upregulated in S3RL and S3DF, respectively (Fig. S7B, Table S5). Analysis of pathway enrichment identified ETC, mitochondrial respiration and mitochondrial translation in tumors from S3RL. Immune responses genes, including genes for major histocompatibility complex II (MHCII), were upregulated in tumors from S3DF patients (Fig. S7C). Notably, the mechanistic connection between altered ETC activity and expression of MHC I genes was recently reported via epigenetic reprogramming (Mangalhara et al., 2023), while expression of MHC II on renal cancer cells has some prognostic values (Saito et al., 1996). We constructed a 23-gene signature containing genes encoding a subset of ETC subunits, mitochondrial ribosomal proteins (MRPs) and MHCII that classified ccRCC for low, intermediate, and high risk of disease recurrence (Fig. 7A). Using the 23 gene signature and leave-one-out (tumor) cross-validation, we developed supervised machine learning models to predict relapse vs. disease free status from gene expression levels. These (Random Forest and LASSO) models achieved high classification accuracy, as indicated by the area under ROC curves (Fig. 7B), supporting strong predictive power of the 23-gene signature.

**Figure 7.**
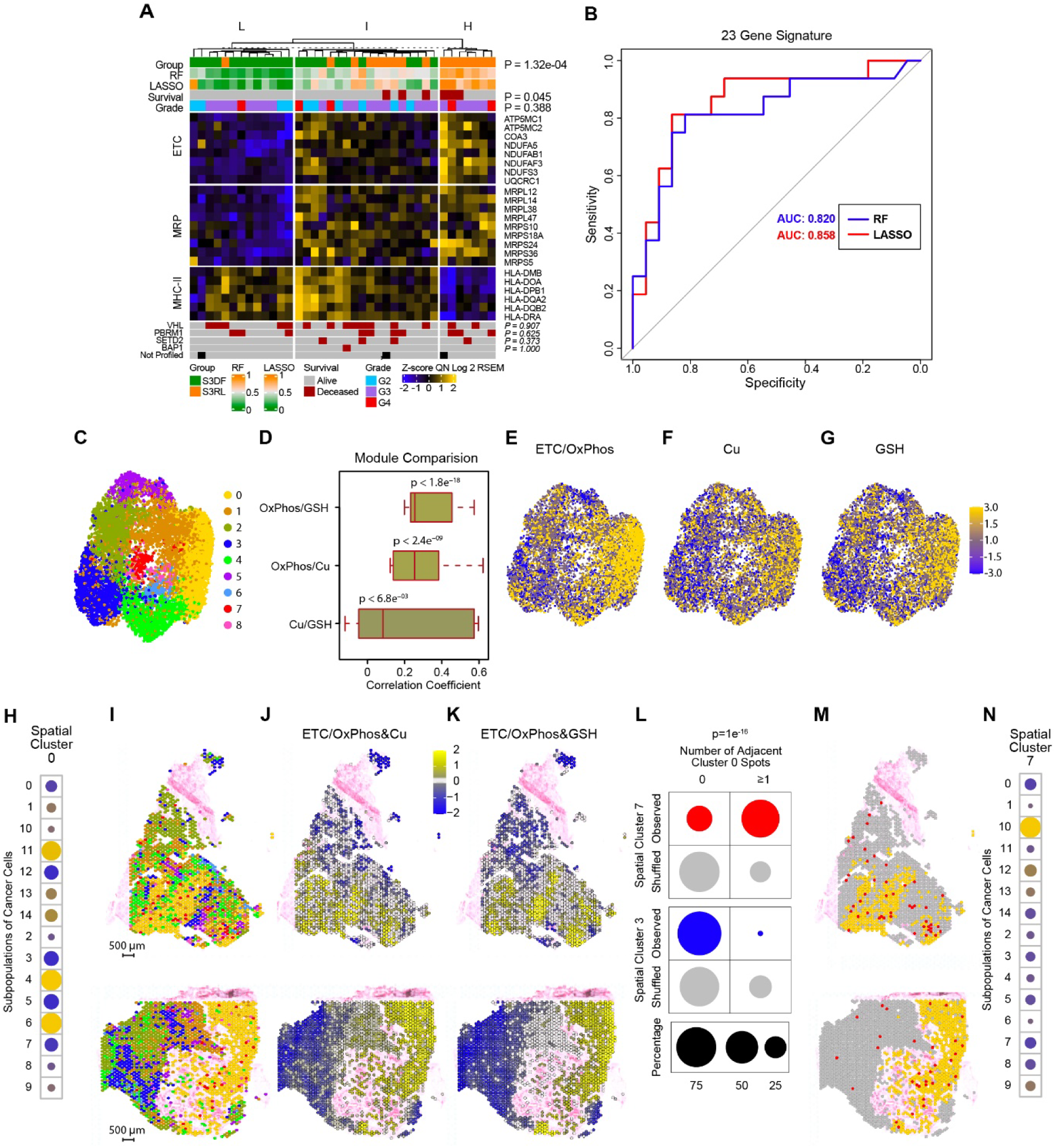
Prognostic application of metabolic signature and adjacent localization of cancer cells with metabolic and proliferative states support the role of metabolic state in tumor progression. (A) Unsupervised k-means clustering (k=3) of tumors from S3DF and S3RL patients using 23 significantly differentially expressed genes belonging to ETC, mitochondrial ribosomal protein (MRP), and MHC class II (MHC-II) gene sets. Samples were stratified into three distinct groups characterized by low (L, n=13), intermediate (I, n=18), and high (H, n=7) risk of disease recurrence, which is further supported by supervised binary classification models, random forest (RF) and penalized regression (LASSO). Fisher’s exact test P values are shown. (B) ROC curves for the classification of the samples into S3DF vs. S3RL classes using random forest (RF) or LASSO classification models. Random classifier is drawn as a diagonal gray line, and classification accuracy is represented by the area under the curve (AUC). (C) UMAP visualization of 9 clusters of spatial spots (spatial clusters) characterized by similar transcriptomic signatures. (D) Pearson correlation of expression of the indicated sets of genes across all 9 spatial spots (E) Annotation of spatial clusters by ETC/OxPhos gene set expression. (F) Annotation of spatial clusters by Cu related gene set expression. (G) Annotation of spatial clusters by GSH related gene set expression. (H) Increased expression of gene signatures of metabolically active scRNA-seq subpopulations 4, 6 and 11 in spatial cluster 0. The color indicates the direction of change in expression of scRNA-seq gene signature for each subpopulation in spatial cluster 0 compared to the mean expression across all spots. Yellow indicates means higher and blue means lower than average expression across all spots. The size of the circles represents the absolute value of the mean of z-score transformed module scores within spatial cluster 0. The one sample t-test for the null hypothesis of mean = 0 for scRNA-seq subpopulations 4, 6 and 11 has p-value < 1e-16. (I) Mapping of spatial clusters into the H&E-stained sections of two stage 3 ccRCCs. Scale bar = 500 µm. Color legend as in panel C. (J) Mapping of concordant expression of ETC/OxPhos and Cu related gene sets into the tumor sections. (K) Mapping of concordant expression of ETC/OxPhos and GSH related gene sets into the tumor sections. (L) Percentage of spots in spatial clusters 7 (shown in red) and 3 (shown in blue) that are adjacent to spots of spatial cluster 0 for observed vs. randomly shuffled (shown in gray) data. The chi-squared test show statistically significant increase in the adjacency between spots of spatial clusters 0 and 7 and decreased adjacency between spots from spatial clusters 0 and 3 in the observed data as compared to the shuffled data. (M) Mapping of spots from spatial cluster 0 (yellow) and 7 (red) on representative sections of ccRCCs. (N) Significant overlap in gene expression between spatial cluster 7 and scRNA-seq subpopulation 10 characterized by proliferative phenotype. The color indicates the direction of change in expression of scRNA-seq gene signature for subpopulation 10 in spatial cluster 7 compared to the mean expression across all spots. Yellow indicates means higher and blue means lower than average expression across all spots. The size of the circles represents the absolute value of the mean of z-score transformed module scores within spatial cluster 0. The one sample t-test for the null hypothesis of mean = 0 for scRNA-seq subpopulation 10 has p-value < 1e-16. The yellow-blue scale shown in panel 7G applies to panels E, F, G, G and N. See also Figure S7 and Table S6.

To reveal spatial organization and local heterogeneity of the cancer cell subpopulations in ccRCCs, we performed 10x Genomics Visium sequencing of five stage 3 ccRCCs from UC Tumor Bank. We identified nine distinct clusters of spots (spatial clusters) based on patterns of differential gene expression (Fig. 7C and Table S6). There was an overall significant positive correlation between expression of ETC/OxPhos, Cu and GSH related gene sets across all spatial clusters (Fig. 7D), which was particularly strong within spatial cluster 0 (Fig. 7E-7G). Moreover, spatial cluster 0 showed significant enrichment for metabolically dynamic single cell subpopulations 4, 6 and 11 (Fig. 7H).

Mapping of the spatial clusters into the tumor sections revealed that in most cases spots from the same cluster were positioned adjacent to one another, consistent with clonal expansion (Figs. 7I, S7D and S7E). Areas of tumors corresponding to spatial cluster 0 showed concordant expression of ETC/OxPhos and Cu related genes (Fig. 7J) or ETC/OxPhos and glutathione related gene sets (Fig. 7K), corroborating scRNA-seq data. In contrast, spots from cluster 7 were scattered across sections.

To assess the adjacency between spatial cluster pairs in tumor sections we quantified the non-randomness in the association of spots from each pair of spatial clusters using observed vs. shuffled spots. Strikingly, we found clear adjacency only between spots from spatial clusters 0 and 7 (Figs. 7L and 7M). Interestingly, spatial cluster 7 had significant similarity to the subpopulation 10 identified in scRNA-seq analysis characterized by proliferative phenotype (Fig. 7N). These results implicate functional interactions between metabolically active and proliferating cells with consequences for tumors progression. In contrast, there was a decrease in adjacency between spatial cluster 0 and 3 relative to reshuffled data (Fig. 7L).

Collectively, the data from three independent cohorts of ccRCCs and comprehensive analysis by single cell, spatial and bulk transcriptomics demonstrate activation of mitochondrial oxidative phosphorylation associated with Cu and glutathione metabolism as a key metabolic process activated during ccRCC progression.

## DISCUSSION

Cu-driven orchestration of metabotranscriptional programs results in oncogenic homeostatic states, which enable ccRCC growth and progression. During evolution of ccRCC there is an increase in Cu levels in the blood and in tumor tissues, leading to increased allocation of Cu to CuCOX and bioenergetic advantages. Reactivation of this respiratory activity, requires, however, adaptive detoxification of harmful reactive oxygen species using a pool of glutathione to prevent toxicity of Cu-H_2_O_2_ Fenton-like reaction and to allow survival of these metabolically dynamic cells. Coupling of glucose utilization for biosynthesis of ATP, generation of ATP by oxidative phosphorylation and biosynthesis of GSH represents an adaptation in redox homeostasis to high Cu. Glucose-derived glutathione has been shown to play role in ROS detoxification in pancreatic islets, although the biochemical pathway required pyruvate carboxylase activity (Fu et al., 2021), while our metabolomic experiments did not show participation of pyruvate carboxylase activity in the biosynthesis of glucose-derived GSH. On the other hand, glutamine metabolism serves as a basic metabolic operating system that is similar in Cu*^Hi^* and Cu*^Lo^* cells. Glutamine-derived GSH supports essential cellular functions such as, for example, protein folding in ER, but that are not specific for Cu*^Hi^*conditions.

We further propose that this Cu-dependent metabolic reprogramming occurs in the subset of cancer cell subpopulations. These metabolically active subpopulations can directly interact with subpopulations of scattered, proliferating cancer cells supporting their rapid growth via released or consumed metabolites. Alternatively, some metabolically dynamic cells may transition to the proliferative state due to their bioenergetic, biosynthetic and redox advantages, or shift to the proliferative subtype due to the accumulation of genetic and epigenetic modifications caused by metabolic reprogramming. Changes in ketoglutarate and succinate have intrinsic and extrinsic epigenetic effects (Dalla Pozza et al., 2020; Liu and Yang, 2021), while ROS affect multiple signaling pathways and genomic stability (Harris and DeNicola, 2020; Yang et al., 2016). Recently, the role of Cu(II) as a direct metal catalyst activating H_2_O_2_-dependent oxidation of NADH to regenerate NAD was proposed to have metabolic consequences for epigenome and macrophage plasticity (Solier et al., 2023).

The mechanisms leading to higher Cu accumulation during ccRCC advancement are currently under investigation. It is possible that exposures of patients to higher levels of environmental Cu, such as tobacco and e-cigarette smoking, result in higher uptake and accumulation in the tumors (Reigle et al., 2021). However, it is also possible that there is a cancer-related organismal level of regulation mobilizing Cu from its liver storage. Positive correlations between Cu levels in tumors and sera, and between total tumor Cu and CuCOX in advanced tumors support increased active uptake of Cu by tumor cells and its selective allocation to CuCOX. The roles of high affinity Cu transporter, CTR1 (Wee et al., 2013), endocytosis, macropinocytosis, an actin-dependent process of nutrient uptake (Aubert et al., 2020), and CD44 (Solier et al., 2023) remain to be determined.

Cu-dependent reprogramming of ccRCC metabolism presents several clinically relevant vulnerabilities. Chelation of Cu has yet to be explored as a therapeutic approach for treatment of ccRCC. However, this may be a double-edged sword. Cu chelation may paradoxically rescue cancer cells with high Cu content in the regions of tumors that are exposed to low glucose concentration leading to resistance. However, the Cu-driven glutathione phenotype of advanced ccRCC may be particularly sensitive to the chemodynamic therapy (CDT), leveraging Fenton-like chemistry to convert intracellular H_2_O_2_ into toxic hydroxyl radicals. Cu is particularly suited to drive this reaction because its efficiency to drive Fenton chemistry is much faster than that of iron and is not affected by pH (Hao et al., 2021). Metal-organic framework (MOF) nanoplatforms are used to deliver Cu and to enhance levels of H_2_O_2_, while their intracellular dissociation utilizes glutathione leading to cell death (Tian et al., 2021). This treatment induces a redox catastrophe similar to that caused by glucose starvation of Cu*^Hi^* cells. Importantly, complementary strategies targeting subpopulations of cells with metabolic and proliferative states should be considered. Finally, measurements of Cu and CuCOX in primary tumors may serve as a simple prognostic biomarker.

Recently, a novel Cu-induced programmed cell death, cuproptosis, was proposed that involves aggregation of lipoylated proteins of TCA cycle, followed by loss of iron-sulfur clusters and proteotoxic stress (Tsvetkov et al., 2022). We did not detect any changes in accumulation, lipoylation or aggregation of lipoylated proteins to indicate the role of cuproptosis in cell death caused by glucose starvation in Cu*^Hi^* cells. This may be related to the different biological context of the chronic, rather than acute, Cu exposure in our experiments. However, engagement of cuproptosis machinery in ccRCC containing high Cu levels maybe an alterantive therapeutic approach.

This study identified a direct role of Cu coordinately driving glucose into full oxidation for biosynthesis of reduced glutathione that is necessary to maintain redox homeostasis. This biochemical and functional compartmentalization of glutathione activity is intriguing, and its mechanism will require further investigation. Glucose-derived GSH limited accumulation of H_2_O_2_ in Cu*^Hi^* cells. It remains to be determined if that activity occurs in mitochondria or cytosol. We did not detect increased expression of known enzymes utilizing GSH to decompose H_2_O_2_, and levels of GPX1/2 were substantially decreased in Cu*^Hi^* cells. It is possible that GSH regulates activity of a novel mechanism leading to removal of H_2_O_2_ or that GSH directly scavenges H_2_O_2_ (Cassier-Chauvat et al., 2023). Interestingly, in vitro, CuCOx binds and can decompose H_2_O_2_ (Jancura et al., 2014), while bacterial cytochrome c peroxidase uses H_2_O_2_ as final acceptor of electrons under anoxic conditions (Khademian and Imlay, 2017). It is possible that similar activities of CuCOX in Cu*^Hi^* cancer cells regulate accumulation of H_2_O_2_ in a GSH-dependent manner. Glucose-derived GSH may be also used by mitochondria to maintain redox state of ETC assembly (Leary et al., 2009; Morgada et al., 2015; Zitare et al., 2015) or contribute to glutathionylation of mitochondrial dehydrogenases and subunits of complex I, preventing formation of superoxide and H_2_O_2_ while maintaining increased OCR (Applegate et al., 2008; Gill et al., 2018; Mailloux et al., 2022; O’Brien et al., 2017).

Interestingly, Cu stimulates pyruvate decarboxylation to acetyl-CoA by inhibiting phosphorylation of PDHA by PDK. Since there was no major change in protein expression of PDK or pyruvate dehydrogenase phosphatase (PDP), it may be the case that their activities are allosterically regulated. PDK is inhibited by pyruvate (Sugden and Holness, 2003). Pyruvate-alanine transfer of cytosolic pyruvate to the mitochondria may result in higher mitochondrial concentration of pyruvate and inhibition of PDK activity. The mechanism of alanine entry into the mitochondria will need to be further investigated. Two transporters are currently known to import small neutral amino acids into mitochondria and may be candidates: SLC25A38 and the members of the sideroflexin family (SFXN). SLC25A38 is a glycine transporter required for heme synthesis (Lunetti et al., 2016). Its gene is located on chromosome 3p and is deleted in 16% of ccRCCs (TCGA KIRC, Firehose Legacy Cohort), consistent with loss of 3p as the most frequent genetic alteration. SFXN1 is a mitochondrial serine importer required for one-carbon metabolism and nucleotide biosynthesis, which has been shown to transport alanine, glycine and cysteine (Kory et al., 2018). It is also critical for ETC integrity and activity (Acoba et al., 2021). *SFXN1* is located on chromosome 5q and is amplified, or its mRNA levels are increased in 29% of ccRCCs (TCGA KIRC, Firehose Legacy Cohort), consistent with 5q amplification as the second most frequent genetic alteration in ccRCC.

Collectively, we demonstrated how organometallic metabolism is deregulated in the process of ccRCC progression and how metabolic plasticity supports a growth and survival advantage of specific clones associated with tumor progression.

### Limitations of the study

Independent cohorts of ccRCCs were used for different measurements. Further validation on a demographically diverse single larger cohort where each tumor can be analyzed by all multiomic studies is needed to strengthen and extend the analysis for potential clinical application. Mechanistically, understanding of the compartmentalization of glutathione pools as well as events by which GSH prevents accumulation of H_2_O_2_ will need to be further investigated. Finally, a fundamental mechanism by which increased Cu exposure leads to the assembly of CuCOX is not known.

## METHODS

All materials are listed in Table S7.

### Data and code availability

The raw RNA sequence data from 786-O cell line were deposited into GEO with accession number <u>GSE250028</u> https://www.ncbi.nlm.nih.gov/geo/query/acc.cgi?acc=GSE250028); the raw sequence data from spatial transcriptomics were deposited into GEO with accession number <u>GSE250163</u> (https://www.ncbi.nlm.nih.gov/geo/query/acc.cgi?acc=GSE250163). Codes were deposited in a public github repository (https://github.com/mjarek66git/ccRCC). Any additional information reasonably needed to reanalyze the data is available from the lead contact.

### Human Specimens

For copper and metabolomic measurements and spatial transcriptomics deidentified ccRCC specimens from white males were obtained from the University of Cincinnati Tumor Bank (IRB exemption 2023-0668). All samples were reviewed by pathologists and derived from region with more than 80% cancer cells. Clinical information regarding tumor recurrence was available through the Tumor Bank. Fresh-frozen tissues were used for metallomic and metabolic assays and FFPE tissues were used for spatial transcriptomics. For scRNA-seq, human data were obtained from two published data sets for 18 primary ccRCCs, labeled as cohort 1 (Li et al., 2022; Obradovic et al., 2021) and cohort 2 (Li et al., 2022). For the TCGA analysis KIRC Firehose Legacy cohort limited to white males was used. Clinical fields from the KIRC dataset were used to define two groups of interest based on patient sex, race, tumor stage, and disease-free survival. ‘S3DF’ consisted of male, white, stage 3 patients who remained disease free for at least 24 months after surgery (n=22). ‘S3RL’ consisted of male, white, stage 3 patients whose disease recurred within 24 months (n=20). Of the 20 patients in the latter group, 4 were removed after oncologist review of pathology reports (available on cBioPortal) due to presence of non-clear cell morphology (sarcomatoid and/or papillary).

### Xenografts

For all experiments 5-week-old Nod-SCID gamma or athymic nude male mice were used. For orthotopic xenografts, mice were administered pre-emptive analgesics. Tumor cells (1×10^6^ in 35 µl volume of 50% Matrigel Corning #354234) were injected unilaterally under the kidney capsule from retroperitoneal approach. The incisions were closed in 2 layers. For subcutaneous xenografts, mice were bilaterally injected with the same number of cells in 200 µl volume of 50% Matrigel. Mouse were divided into two groups, one fed with Cu deficient diet (Envigo Teklad TD.80388), and the other with either standard chow or matched custom-made high Cu diet (Envigo Teklad, TD.220421). In the case of subcutaneous tumors, tumor’s dimensions were measured weekly using calipers and tumor volume was calculated using formula 1/2xWxWxL.

### Cell lines

Human ccRCC cell lines 786-O and RCC4 were carried in DMEM/F12 (Cytiva #SH30023) + 10% fetal bovine serum (Gibco #16000-044) at 37°C and 5% CO_2_. Cell lines were regularly authenticated by STR profiling (LabCorp, Burlington, NC) and tested for mycoplasma (MycoAlert, Lonza #LT07-218). To create a chronic Cu*^Hi^*exposure model, cells were adapted to high Cu conditions by gradually increasing Cu concentration by switching them over a two-week period to media containing 7.5, 15, 22.5 and 30 µM Cu, and then were carried in DMEM/F12 with 10% FBS medium containing 30 µM Cu (Fig. S1K). To generate high Cu media that models Cu bound to blood proteins, 100% FBS was first incubated overnight with CuSO_4_ at 4°C, and then ten-fold diluted with the media. Binding of Cu to serum proteins and final total concentration of Cu in the media was routinely validated by ICP-MS. Cells were used up to 15 passages from the switch to Cu*^Hi^* media.

### Metallomics

Total element and element distribution of tumors or cell lines lysed were analyzed by ICP-MS and SEC-ICP-MS as described in detail in (Reigle et al., 2021). The total Cu concentration was determined by the external calibration method in an Agilent 8900 ICP-MS/MS system. For quality control analysis, a certified reference material (NIST® SRM® 1577c) was analyzed in parallel, with sulfur used to normalize the Cu content. The size exclusion separations were carried out in a TSKgel QC-PAK GFC 300 column (7.8 x 150 mm, 5µm) in an Agilent 1260 Agilent HPLC equipped with a quaternary pump, an autosampler and a diode arrange detector (DAD). This analysis was performed with a mobile phase composition of 50 mM ammonium acetate in 0.5% methanol. The flow rate of the mobile phase was 0.65 mL min^-1^ and the injection volume was 80 µl. The LC captured the full UV-Vis spectra from 220 to 750 nm, while the ICP-MS/MS system was operated in TRA mode, under helium collision mode (at 3 mL/min), with integration time of 0.1 s including ^63^Cu, ^65^Cu and ^34^S, isotopes. The quantification was performed by the external calibration method, after integrating the peak areas of the Cu chromatograms of a mix of metalloprotein standards (GFS, 1511901, Bio-Rad Laboratories, Inc.) and normalized by the total area of the respective chromatogram at 280 nm.

### Metabolomics

For glucose and glutamine tracing experiments, 786-O cells were plated in quadruplicate in 60 mm plates in DMEM/F12 (without glutamine and glucose) (Biowest #L0091) supplemented with 10 mM glucose, 2.5 mM glutamine, and 10% dialyzed FBS (Gibco 26400-044). After 24 hours media was changed to the same media supplemented with 5 mM glucose (unlabeled or ^13^C_6_ labeled), and 2.5 mM glutamine (unlabeled or ^13^C_5_,^15^N_2_ labeled) and 10% dialyzed FBS and collected at the indicated time points. For pyruvate tracing experiment, cells were changed to DMEM (Gibco A1443001) supplemented with 0.05 mM glucose, 4 mM glutamine, 2 mM ^13^C_2_ pyruvate, and 10% dialyzed FBS, with or without 10 µM BSO for 16 h. For each time point, cell number on an additional plate was counted for normalization. Media from each sample was collected and stored at -80 °C for analysis. For each time point, cells on an additional plate were harvested and counted for normalization. Remaining cell plates were washed 3x with 2 mL of cold PBS. Following the final wash, PBS was aspirated well, plates were placed onto dry ice, then 0.9 mL of ice cold 5:3:2 MeOH:MeCN:water (v/v/v) was added to each. Plates were scraped and lysate transferred to pre-chilled 1.5 mL microcentrifuge tubes. Metabolomics extractions and analyses were performed as previously described (Nemkov et al., 2019). Extracts vortexed for 30 min at 4°C, clarified by centrifugation (10 min, 15,000 g, 4°C), then 500 µL of supernatant dried under vacuum. Samples were shipped as dry residue to the University of Colorado School of Medicine on dry ice. Residue was reconstituted in 5:3:2 MeOH:MeCN:water (v/v/v) according to cell counts to a normalized concentration of 2e6 cells/mL. The metabolite extracts were analyzed (10 µL per injection) by ultra-high-pressure liquid chromatography coupled to mass spectrometry (UHPLC-MS -Vanquish and Q Exactive, Thermo). Metabolites were resolved on a Phenomenex Kinetex C18 column (2.1 x 150 mm, 1.7 µm) at 45°C using a 5-minute gradient method in positive and negative ion modes (separate runs) over the scan range 65-975 m/z exactly as previously described (Nemkov et al., 2019). Following data acquisition, .raw files were converted to .mzXML using RawConverter then metabolites assigned and peaks integrated using Maven (Princeton University) in conjunction with the KEGG database and an in-house standard library. ^13^C_2_ isotopologue peak areas were corrected for the natural abundance of ^13^C_2_ in the parent compound. Quality control was assessed using technical replicates run at beginning, end, and middle of each sequence as previously described (Nemkov et al., 2015; Nemkov et al., 2017).

### Lipidomics

Lipids were extracted from purified mitochondria or xenograft samples (10 mg/sample) using a modified Bligh and Dyer method. Briefly, mitochondrial pellets or xenograft tissues were homogenized in water:methanol:dichloromethanane (DCM) 1:1:1 (v/v/v) and centrifuged at 2671 g for 5 min. The upper layer (organic) was collected, and the bottom layer (aqueous) was re-extracted by phase separation. The organic layers were combined and dried using a SpeedVac Concentrator. Extracts were resuspended in DCM/methanol/isopropanol (2:1:1, v/v/v) containing 8 mM ammonium fluoride. SPLASH LipidoMix™ was used as internal standard. A bicinchoninic acid (BCA) protein assay was performed on the aqueous protein layer to normalize the lipid concentrations in each sample.

Reversed-phase chromatographic separation was achieved with the Accucore C30 column: 3 μm, 2.1×150 mm (Thermo Fisher Scientific, Waltham, MA). The column was maintained at 35°C and tray at 20°C. Solvent A was 10 mM ammonium formate (AF, LC-MS grade) in 60:40 Acetonitrile (ACN):water (LC-MS grade) with 0.1% formic acid (FA, LC-MS grade). Solvent B was composed of 10 mM AF with 90:10 IPA:ACN with 0.1% FA. The flow rate was 250 μl/min, and the injection volume was 10 μl. The gradient was 50% solvent A (3 to 50%). The Orbitrap (Thermo) mass spectrometer was operated under heated electrospray ionization (HESI) in positive and negative modes separately for each sample. The spray voltage was 3.5 and 2.4 kV for positive and negative mode, the heated capillary was held at 35°C and heater at 27°C. The sheath gas flow rate was 45 units, and auxiliary gas was 8 units. Full scan (m/z 250–2000) used resolution 30,000 at m/z 200 with automatic gain control (AGC) target of 2×10^5^ ions and maximum ion injection time (IT) of 100 ms. Normalized collision energy (NCE) settings were 25%, 30%, 35%.

Lipid identification and relative quantification were performed with LipidSearch 4.1 (Thermo) (Breitkopf et al., 2017), followed by Lipidsig (Lin et al., 2021). The search criteria were as follows: product search; parent m/z tolerance 5 ppm; product m/z tolerance 10 ppm; product ion intensity threshold 1%; filters: top rank, main isomer peak, FA priority; quantification: m/z tolerance 5 ppm, retention time tolerance 1 min. The following adducts were allowed in positive mode: +H, +NH_4,_ +H H_2_O, +H 2H_2_O, +2H, and negative mode: -H, +HCOO, +CH_3_COO, -2H. Cardiolipins were identified as -H, -2H and +HCOO adducts.

### Oxygen consumption rate

OCR was measured using a Seahorse XFe96 Analyzer (Agilent #S7800B). 48 h after plating media was changed to DMEM (Agilent #103575-100) + 10 mM glucose + 4 mM glutamine + 0.5 mM pyruvate or other indicated condition and incubated in a 37°C non-CO_2_ incubator for 1 h before assay. OCR was measured after each sequential injection of 1 µM Oligomycin A, 0.25 µM (786-O) or 2 µM (RCC4) FCCP, and 0.5 µM Rotenone/0.5 µM Antimycin A/2.5 µM Hoechst. To calculate basal (pre-drug injection), ATP coupled (drop post Oligomycin A injection), and maximal (post FCCP injection) OCR rates, data was exported to a Seahorse XF Cell Mito Stress Test Report Generator using the Seahorse Wave Desktop Software (Agilent). Hoechst fluorescent intensity was used to normalize for cell number.

### Organelles isolation and fractions

Mitochondria isolation was performed using the Miltenyi Mitochondria Isolation Kit, human (#130-094-532). Cells were pelleted and resuspended in 500 µL lysis buffer supplemented with 1x protease and phosphatase inhibitor cocktail (PPIC, Thermo Scientific 1861281) and 10 µM phenylmethylsulfonyl fluoride (PMSF, Sigma #P7626, CAS# 329-98-6). Cells were lysed by passing 13 times through a 27G needle. Equal amounts of lysates were diluted with 4.5 mL of 1x separation buffer and rocked at 4°C for 1 h with 50 µL anti-TOM22 microbeads. LS columns were pre-washed, loaded with samples and washed 3 times with 3 mL separation buffer. Beads were eluted by removing columns from magnet and plunging with 1 mL separation buffer and centrifuged at 15,000 g for 15 minutes.

For mitochondrial enriched fraction, cells were lysed as described above. Nuclei were pelleted at 800 x g for 5 min. Mitochondria were pelleted at 15,000 x g for 15 min, washed once with PBS and lysed in RIPA buffer.

For nuclear soluble fraction, cell pellets were incubated 5 min on ice in 10 mM HEPES pH 7.5, 10 mM KCl, 1.5 mM MgCl_2_, 1 mM DTT, 10% glycerol, 0.1% Triton X-100, and 1X PPIC. Samples were centrifuged at 1700 x g for 5 min and supernatant taken as the cytoplasmic fraction. Nuclear pellets were extracted in buffer containing 10 mM HEPES pH 7.5, 300 mM NaCl, 1.5 mM MgCl_2_, 1 mM DTT, 10% glycerol at 4°C for 1 h. Samples were centrifuged, and the supernatant collected as the soluble nuclear fraction.

### SDS-PAGE and immunoblotting

Samples were run on Bio-Rad Criterion XT 12% (#345008) or 4-12% (3450124) Bis-Tris gels with 1x NuPAGE MOPS running buffer (Thermo #NP0001). Protein was transferred to 0.2 µm PVDF membranes. Antibodies are listed in the Key Resources Table. Blots were developed using Pierce ECL Western Blotting Substrate (Thermo #32106), SuperSignal West Femto Maximum Sensitivity Substrate (Thermo #34095), or SuperSignal West Atto Ultimate Sensitivity Substrate (Thermo A38555) and imaged on a Bio-Rad ChemiDoc Imaging System. Restore PLUS Western Blot Stripping Buffer (Thermo #46430) was used to strip blots for re-probing. Blots or scanned gels were quantified using Image Studio Lite software (LiCor) or ImageJ.

### Blue Native-PAGE

Mitochondria were extracted with 1x sample buffer (NativePAGE Sample Prep Kit, Invitrogen #: BN2008) containing 1% digitonin for 15 minutes on ice. Lysates were cleared by centrifuging at 20,000 g for 20 minutes, then 0.25% G-250 Sample Additive was added to supernatant. Samples were run on NativePAGE 3 to 12% Bis-Tris gels (Invitrogen #BN1001BOX) with NativeMark Unstained Protein Standard (Invitrogen #LC0725). Running buffers were prepared using the NativePAGE Running Buffer Kit (Invitrogen #BN2007). 1x running buffer was used as the anode buffer in a Thermo XCell SureLock Mini-Cell (#EI0001). Gels were run at 150 V for 40 min with dark blue cathode buffer (1 x cathode buffer additive in running buffer), then ran for 2 h using light blue cathode buffer (0.1 x cathode buffer additive). Gels were transferred to 0.2 µm PVDF for 1.5 h at 40 V in transfer buffer containing 25.6 mM Tris and 190.55 mM glycine. After transfer, the PVDF was fixed in 8% acetic acid for 15 minutes, destained several times with methanol, and used for western blot analysis.

### Respiratory Complex IV In-gel Activity Assay

Mitochondria lysates were prepared and run on a gel as described above for BN-PAGE with the following modification: light blue cathode buffer was used to run the gel for the first 40 minutes and was then switched to clear cathode buffer (no cathode buffer additive) for the remaining running time. After running, gels were rinsed several times with water and incubated an hour-overnight at 37°C in 45 mM phosphate buffer pH 7.4, 0.5 mg/mL diaminobenzidine, and 1mM bovine cytochrome c. After band development, gels were rinsed with water and scanned.

### Cytotoxicity measurements

Cytotoxicity was measured by positive staining with propidium iodide (PI) and/or Annexin V by flow cytometry using the Invitrogen FITC Annexin V/Dead Cell Apoptosis Kit (#V13242). Cells were plated, and 24 h (RCC4) or 48 h (786-O) h later media was changed to DMEM + 10% FBS + indicated reagents. The analysis was performed 72 h (RCC4) or 24 h (786-O) later. Cells were trypsinized, pelleted, and resuspended in 1X annexin binding buffer with 1 µg/mL PI and 1:20 FITC Annexin V. Fluorescence for 20,000 events/sample was measured with a 488 nm laser and 505LP/530/30 emission filter (FITC) and a 561 nm laser with a 600LP/610/20 emission filter (PI) on a BD LSRFortessa.

### BSO Dose Response Curves

Drug sensitivity dose curves were generated using the CyQUANT NF Cell Proliferation Assay (Invitrogen C35006) using 96 well plates. BSO was added 1:1 to cell plate in six technical replicates. After 48 h, media was aspirated and 30 µL of 1x HBSS + 1:1000 dye + 1:1000 dye delivery reagent was added to each well. Plates were incubated for 30 min at 37°C and 5% CO_2_. Fluorescence was measured with excitation at 485 nm and emission at 530 nm on a BMG Labtech CLARIOstar. Fluorescence was normalized relative to control.

### RNAi approaches

RCC cell lines were transfected with siRNAs at final concentrations of 50 nM using Lipofectamine 3000 according to manufacturer’s protocol. Viral transductions included addition of 2 µg/mL polybrene (Millipore TR-1003-G) before viruses were administered. All lentiviral shRNA constructs were VSV-G envelope packaged, concentrated, and used at a 1:40 dilution. All controls were treated with empty, non-target, or scrambled constructs.

### Glutathione measurements

Glutathione was measured using the GSH-Glo Assay from Promega (V6911) according to the manufacturer protocol. 786-O and RCC4 cells were plated in duplicates on a 96 well white-walled plate. Media was changed to DMEM + 10% FBS with indicated glucose or glutamine concentrations. For glutathione measurements cells were incubated in 50 µL of 1x GSH Glo Reaction Buffer + 1:100 Luciferin Substrate + 1:100 Glutathione-S-transferase (+1mM DTT for GSH + GSSG measurement) for 30 min at room temperature. 50 µL of reconstituted Luciferin Detection Reagent was added and plate was incubated another 15 min at room temperature. Luminescence was measured on a BMG Labtech CLARIOstar. A negative control was subtracted from each value. To calculate GSSG measurement, a non-reduced (GSH) sample measurement was subtracted from a reduced sample (GSH + GSSG) measurement. A matched plate was treated with the Cyquant NF Proliferation assay reagent and fluorescence used to normalize glutathione measurement to cell count.

### ROS measurements

Cells were plated and treated as described in the *Cytotoxicity* section. Cells were treated with 500 nM MitoSOX Red (Invitrogen M36008) and 100 nM MitoTracker Green (Invitrogen M7514) or 1.5 µM CM-H2DCFDA (Invitrogen C6827) for 30 min at 37°C, washed, trypisinized, and pelleted, then resuspended in PBS + 1% BSA + 0.1 µg/mL DAPI (Invitrogen D3571). The geometric mean of the fluorescent intensity was measured with a 488 nm laser and 505LP/530/30 emission filter (MitoTracker Green/CM-H2DCFDA), 405 nm laser and 600LP/610/20 emission filter (MitoSOX red), and 405 nm laser and 450/50 emission filter (DAPI) on a BD LSRFortessa for 20,000 events/sample. DAPI was used to gate for live cells. An unstained control was used to subtract background fluorescence.

### Histology

Tumor sections were stained with hematoxilin and eosin (H&E) or analyzed by immunohistochemistry performed in Pathology Research Core at CCHMC according to standard protocols.

### Immunofluorescences

Cells plated on glass coverslips were fixed with 4% paraformaldehyde for 20 min at room temperature or 100% methanol for 5 min at -20°C. Cells were permeabilized with 0.1% saponin, blocked with PBS containing 0.1% saponin and 1% BSA for 30 min and incubated with primary antibody for 1 h at 37°C. Coverslips were then washed and incubated with Alexa Fluor labeled secondary antibodies for 30 min at room temperature. Finally, coverslips were washed and mounted using DAPI Fluoromount-G and analyzed by confocal microscope. Antibodies and concentrations are listed in the STAR Table.

### RT-PCR

RNA was purified from cells using TriReagent (MRC #TR 118) according to manufacturer protocol, resuspended in nuclease free water, and its concentration was determined with a NanoDrop. cDNA was synthesized using the Applied Biosystems High-Capacity cDNA Reverse Transcription Kit (#4368814). Final cDNA was diluted to 8 ng/uL. qPCR was performed using 1X Fast SYBR Green Master mix (Applied Biosystems #4385612), 400 nM primers, and 20 ng cDNA using the Fast setting on an Applied Biosystems QuantStudio7. Primers are listed in the STAR Table. Analysis was performed using the ΔΔCt method using PP1A as the housekeeping gene and normalizing relative to Cu*^Lo^*.

### Statistical analysis

Data points represent biological replicates and data are expressed as mean ± SEM. Significance was determined using two-tailed t-test, one sample t-test, paired t-test or one-way or two-way ANOVA with Holm-Sidak post-hoc test as indicated using SigmaPlot v14/v15 or GraphPad Prism. Graphs were created in GraphPad Prism.

### RNA-seq

RNA was extracted using RNAlater ICE (Ambion) and miRNA isolation kit (Ambion). The quality of RNA was checked using Bioanalyzer RNA 6000 Nano kit (Agilent). PolyA RNA was extracted using NEBNext Poly(A) mRNA Magnetic Isolation Module (NEB) and used as input for RNA-seq. RNA-seq was performed in UC Sequencing Core. RNA-seq libraries were prepared using NEBNext Ultra II Directional RNA Library Prep Kit (NEB). After library QC analysis using Bioanalyzer High Sensitivity DNA kit (Agilent) and library quantification using NEBNext Library Quant kit (NEB), the sequencing was performed under the setting of single read 1×51 bp to generate ∼30 million reads per sample on HiSeq 1000 sequencer (Illumina).

Differential gene expression analysis was performed using *edgeR*, comparing three Cu*^Lo^* and Cu*^Hi^* samples. Gene Set Enrichment Analysis (GSEA) was conducted using the *clusterProfiler* package (Wu et al., 2021) in R, utilizing a curated library that integrates an in-house NRF2 signature with KEGG terms. The input ranked gene list for GSEA was obtained by the *edgeR* generated log2 fold-change in the descending order. Subsequently, the *dotplot* and *gseaplot* functions from the *clusterProfiler* package were employed to generate the enrichment and GSEA plots. The top 200 differentially expressed genes (DEGs) identified by *edgeR* were subsequently used to stratify TCGA ccRCC cohort. Specifically, TCGA_KIRC_RNASeqV2_2019 dataset (n=309) from *ilincs* portal and K-means (k=2) algorithm were used to perform clustering analysis for both genes and tumor samples, limited to white males only (Fig. 1S). Consensus pathways that were enriched in both 786-O Cu*^Hi^* cells and in the subpopulation 6 identified in scRNA-seq were ranked using Fisher consensus of adjusted p-values obtained from individual enrichment analyses (Fig. 1U).

### scRNA-seq analysis

scRNA-seq data for 18 ccRCCs were acquired from two published studies: Cohort 1 (Obradovic et al., 2021) and Cohort 2 (Li et al., 2022). Cells were excluded if they had fewer than 1,000 detected UMIs, expressed fewer than 500 genes, or had mitochondrial gene content greater than 10%. Genes expressed in fewer than 20 cells were also discarded. *Harmony* (Korsunsky et al., 2019) integration algorithm was used in conjunction with *Seurat V4* standard preprocessing pipeline to integrate single cell datasets from different tumors while removing confounding batch effects. Thirty principal components and Louvain algorithm were used to perform dimensionality reduction and clustering in *Seurat*, using *RunPCA*, *RunUMAP*, *FindNeighbors*, *FindClusters* functions. Cancer cells were identified using expression of CA9 and predicted loss of chromosome 3p. Copy number inference was conducted using the *inferCNV* R package (Tickle et al., 2019). Proximal tubular cells, the origin of ccRCC cells (Young et al., 2018), identified in both cohorts via label transferring (Stuart et al., 2019) were used as the reference cells for *inferCNV* analysis. U-maps for visualization of cancer cell heterogeneity were generated using *Seurat*. The expressions of gene sets were computed using *Seurat AddModuleScore* function, employing 24 bins and 100 random genes as the background for each bin.

To assess cluster stability, a subsampling was performed by randomly selecting 80% of the cells from the original ccRCC single cell datasets for 100 times. Jaccard Index was computed as a measure of cluster stability by comparing clusters obtained from the original dataset with those obtained using 100 subsamples (Tang et al., 2021), and plotted using *ggplo2* in R.

The residual contributions to the Pearson correlation coefficient (PCC) were used to assess patterns of correlations between gene sets expression levels across subpopulations. To define the PCC residuals, the covariance of the two gene sets was computed using cells from each subpopulation and divided by the product of their standard deviations calculated using cells from all cells.

### Visualization of metabolic and other gene sets expression patterns

An in-house developed heatmap function (https://github.com/mjarek66git/ccRCC) was used to generate the heatmap in Fig. 6E. Seurat normalized and batch corrected gene expression values for the top 3,000 most variable genes were quantile re-normalized per cell and the distributions were shifted by a constant to obtain all positive values. Cells with less than 500 counts and genes with less than 100 counts were excluded. Hallmark, KEGG and curated metabolic genes sets were used. The expression of the gene set was defined as the average of individual gene expression values. For visualization, gene set expression values across all cells were discretized, with the values in the top quartile (high expression) shown as yellow, the bottom quartile (low expression) as blue, and intermediate values as black. Gene sets (columns) were clustered using hierarchical clustering and 1-PCC as the dissimilarity measure, while cells (rows) were ordered using scRNA-seq cancer cell subpopulations form the scRNA-seq *Seurat* analysis. The order of cell clusters was defined using hierarchical clustering of cluster centroids and 1-PCC as the dissimilarity measure. Uninformative gene sets that did not show association with the Seurat cell clusters or did not cluster with at least one other supergene at the level of PCC=0.5 (note the built-in redundancy between KEGG, Hallmark and curated supergenes) were excluded from the heatmap.

### Bulk RNA data analyses

TCGA Firehose Legacy Kidney Renal Clear Cell Carcinoma data was used to analyze differential gene expression between S3DF and S3RL tumors. The ‘RNA_Seq_v2_expression_median’ dataset downloaded from *cBioPortal*, consisting of RSEM values for 20,531 genes across 534 samples, was transformed using log2(RSEM+1) and quantile re-normalized column-(sample-)wise. Z-score transformation was applied to gene expression rows to emphasize the relative expression value of a gene across samples. Differential gene expression between S3DF and S3RL tumors was assessed using the *t.test* package.

### Significant differentially expressed genes heatmap

*ComplexHeatmap* was used to generate a gene expression heatmap for 1,267 differentially expressed genes with t-test p-values <= 0.05. Samples were partitioned into two clusters (k=2) using *kmeans*. Hierarchical clustering was performed on both rows and columns (within each k-means cluster) using Euclidean distance and average agglomeration. Fisher’s exact test was performed with *fisher.test* to determine the significance of sample group separation between k-means clusters, and other classes of samples defined by tumor grade, survival, and mutation type.

### Gene Set Enrichment Analysis (GSEA)

GSEA v4.1.0 software was used to assess pathway enrichment between groups, using the MSigDB c5.all.v7.2 collection, consisting of 14,765 ontology gene sets. Signed fold change values, scaled by the corresponding p-values from the earlier differential expression analysis, i.e., -log10(p-value)*sign(log2foldchange), were used to rank genes.

### Signature heatmap

Of the 1,267 genes identified to be differentially expressed, electron transport chain (ETC), mitochondrial ribosomal protein (MRP), and MHC class II (MHC-II) genes were combined to generate a 23-gene signature capable of stratifying the samples into distinct clusters. To assess the discriminatory power of such defined signature, samples were partitioned into three slices (k=3) using *kmeans,* while hierarchical clustering was performed on columns (within each cluster) using Euclidean distance and average agglomeration. Fisher’s exact test was performed with *fisher.test* to determine the significance of sample group separation between clusters, and *ComplexHeatmap* was used to generate the heatmap.

### Supervised classification

S3DF vs. S3R class membership was predicted for each sample using random forest (RF) machine learning models trained and assessed using one-sample-leave-one-out cross-validation framework and 23 gene expression values as input features. R function *randomForest* was called with default parameters for training and the class label for the left-out sample was predicted using *predict.randomForest* based on the RF majority vote. For each of the 38 RFs, feature (gene) importance was assessed by mean decrease in Gini index. Additionally, the proportion of trees voting S3RL was recorded for use in later ROC curve generation. In addition, penalized LASSO regression models were trained and assessed using the same cross-validation framework using *cv.glmnet* function.

### Gene correlation density plots

R packages *overlapping* and *ggplot2* were used to generate density plots of pairwise gene correlations. Spearman correlation coefficients (SCC) were calculated for all pairs in the group of 145 metabolic genes in S3DF and S3Rec separately. Gene pairs including genes from a gene set of interest were subsetted and the SCC of those pairs were plotted as density functions. Vertical dashed lines were placed per visual inspection to inform the selection of SCC cutoff for circos plot visualization.

### Spatial transcriptomics

The experiments were carried out in the Advanced Genomics Core at the University of Michigan according to Visium 10x protocol (10xGenomics). FFPE sections of tumor fragments were analyzed by the pathologist and regions containing predominantly cancer tissues with RNA quality higher than DV200 50% were used as capture areas, H&E stained and analyzed for spatial transcriptomics. The results were processed by spaceRanger software to acquire expression matrix of each sample. Seurat-V4 software was used for further processing and data normalization. As a quality control step, spots with fewer than 500 UMIs or more than 30% mitochondrial or ribosomal reads were filtered out.

### Identifying cancer cell spots in the spatial slide and cancer specific clusters

To analyze the cancer cell regions, spatial spots from each slide were clustered using Seurat-V4 and the pipeline used to analyze scRNA-seq data. Small spatial clusters with very low total mRNA counts, likely corresponding to connective tissue were removed. To further discard regions of slides with high stromal or immune cells, we applied the label transfer in Seurat (Stuart et al., 2019) and the distinct cell type expression profiles identified in scRNA-seq analysis from (Young et al., 2018) cohort. Finally, the selected regions were verified using CA9 expression.

### Joint clustering of cancer spots from the five ccRCCs

Normalized gene expression profiles of cancer cell were integrated and clusterd using Seurat-V4. The integration features were selected from the 2000 most variable genes using SelectIntegrationFeatures, and the integration was performed by using PrepSCTIntegration, FindIntegrationAnchors and IntegrateData functions. The Louvain algorithm was applied on 30 principal components using FindNeighbors and FindClusters functions. To evaluate the optimum number of spatial clusters FindClusters function was performed with resolution values ranging from 0.3 to 1.2 at intervals of 0.05, and cluster quality was evaluated at each resolution. For each resolution value, the clustered cells were subsampled 100 times with 500 spots, and silhouette score was computed for these 500 spots and their cluster labels and Pearson correlation was used as the distance metric in computation of silhouette score in the silhouette function. This procedure was repeated for 100 random samples of 500 spots to compute a mean and standard deviation of average silhouette score at each resolution value. The highest resolution value that maximizes mean silhouette score before a major reduction in the silhouette score was selected as the optimal resolution. Differential Gene Expression between clusters was computed by the MAST hurdle model for single-cell gene expression modeling, as implemented in the Seurat FindAllMarkers command, with log fold change threshold of 0.25 and minimum fractional expression threshold of 0.25, indicating that the resulting gene markers for each cluster are restricted to those with log fold change greater than 0.25 and non-zero expression in at least 25% of the cells in the cluster.

### Gene set module score for ETC/OxPhos, Cu and GSH and their association with spatial clusters

The module score of metabolic gene modules were calculated for each cancer spot using Seurat-V4 AddModuleScore function with 24 bins and 100 control genes. The association of these module expressions with the spatial clusters was analyzed by correlating discretized values of module scores with spatial clusters. Module scores were discretized using three bins defined as the 1 quartile, between the 1 and 3 quartile and more the 4 quartile, thus defining 3 states with low, medium and high expression of ETC/OxPhos, Cu and GSH gene modules as shown in (Fig.7).

### Finding association between the single cell subpopulations of cancer cells and the spatial clusters

The module score of top 100 differentially expressed genes in each single cell subpopulation was calculated for the cancer cell spots in the tumor section using AddModuleScore function in Seurat-V4. The module scores were then standardized using scale function in R. For each spatial cluster, the null hypothesis of mean of zero for each normalized module score of single cell clusters’ gene sets was tested using one sample Student’s t-test using t.test function in R. The P value and deviation from the mean of zero is reported for each spatial/single cell cluster pair.

### Spatial organization of spot clusters

For each cluster, we generated normalized distributions of the number of adjacent spots from the same cluster. To assess non-randomness of the distribution, spot labels were re-shuffled while keeping the same number of spots in each cluster. The adjacency of spots from the same cluster was calculated as the difference in the distribution between the observed and shuffled data using Kolmogorov-Smirnov test. To quantify adjacency between a reference cluster (eg. cluster 0) and other clusters, the Chi-squared test was calculated by comparing the number of reference cluster spots with zero vs. more than one adjacent spot from other clusters using observed vs. reshuffled spot labels for the other clusters.

## Acknowledgments

The following grants supported the work: MCK (R01GM128216, R01CA287260 and 2I01BX001110 BLR&D VA Merit Award); JTC (R01CA230904 and R35 GM133561); DRP (R01CA239657); PPS (R01CA259845 and LCS Foundation); KEV (R35GM146878); KCP (R37CA272854); SG (K08CA273542); MEB (T32CA17846); BS (T32CA236764); DS (T32ES007250); JAR, and ADA, on behalf of the University of Colorado SOM Metabolomics Core, acknowledge support from the University of Colorado Cancer Center via NCI P30CA046934 and technical contribution of Rachel Culp-Hill.

Graphical abstract was generated using biorender.com (agreement number GU26CG2W3R). We thank Birgit Ehmer for preparation of immunofluorescent images; Rose Bacon and Lindsey Huether for preparation of the figures.

## Authors Contributions

**Conceptualization:** MCK, JTC, JM, JALF, DRP, PPS, SG, MEB, BS, BV, JAR, ADA, KV, KCP

### Data Curation

**Formal Analysis:** MEB, JY, BS

**Funding acquisition:** MCK, SG, PPS, JALF, JTC, DRP, ADA, KCP

**Investigation:** MEB, JY, BS, DS, BV, RCH, JAR, CB, AM, JB

**Methodology:** MEB, JY, BS, BV

**Project administration:** MCK

**Resources:** BS, JM, RA

### Software

**Supervision:** MCK, JM, JALF, JTC, ADA, PPS

### Validation

**Visualization:** MEB, JY, BS, DS, BV, JAR, CB, AM, RA

**Writing-original draft:** MCK, JM, MEB

**Writing-review and editing:** All authors

**Declaration of Interests:** All authors report no conflict of interest.

---

**Figure S1.**
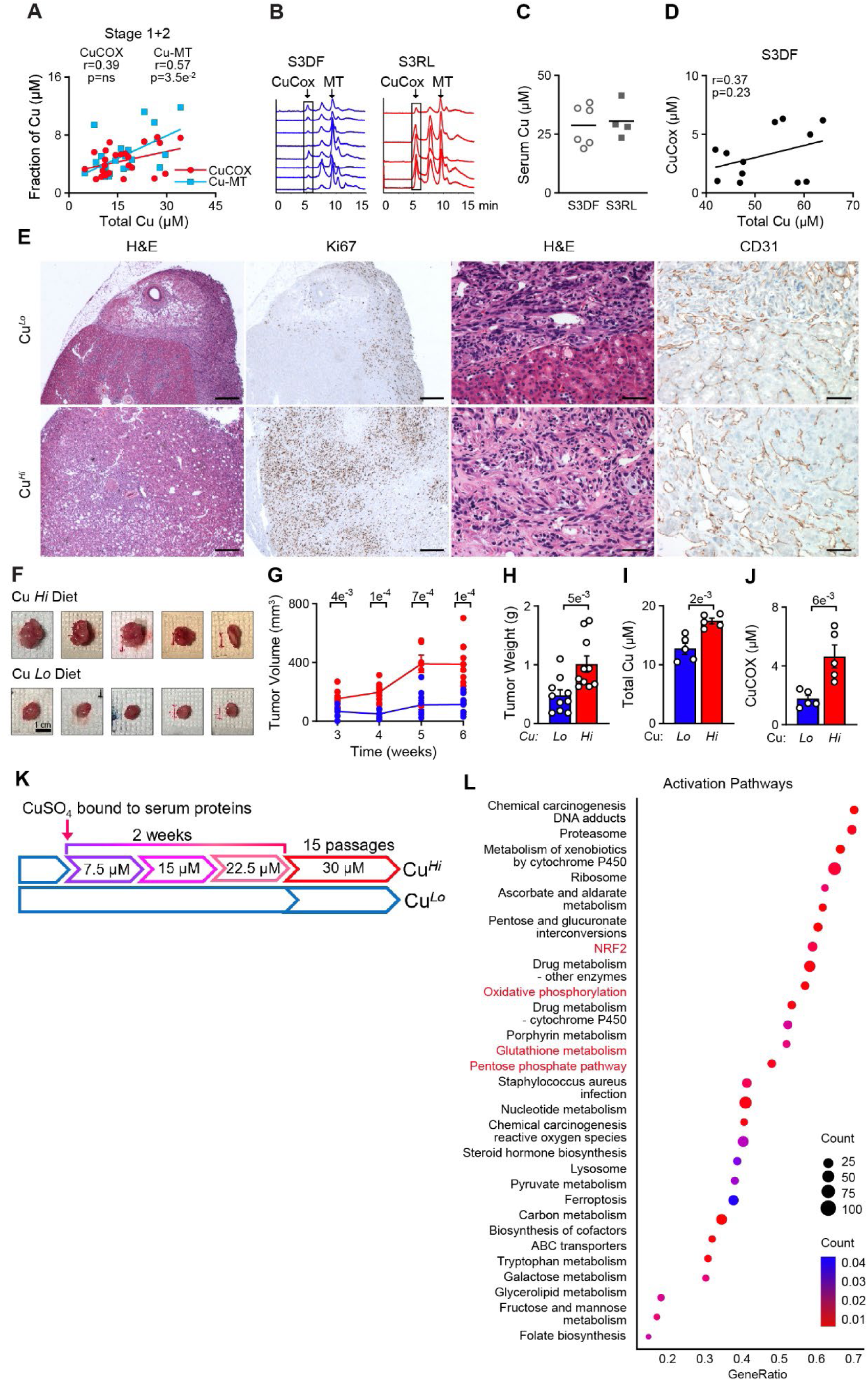
**related to Figure 1. Accumulation of Cu and its allocation to CuCOX associates with progression of ccRCC and stimulates tumor growth.** (A) Pearson correlation of total tumor Cu levels with MT-Cu but not Cu-COX in tumors from stage 1+2 patients. (B) Representative chromatograms from SEC-ICP-MS from S3DF and S3RL tumors; MT-metallothioneins. (C) Levels of Cu in sera from S3DF (n=6) and S3RL (n=4) patients. (D) Pearson correlation of total tumor Cu levels with Cu-COX in tumors from S3DFpatients. (E) Staining for H&E and IHC of proliferation marker Ki67 and of blood vessel marker CD31 in sections from orthotopic xenografts tumors from mice fed matched low and high Cu diet. Scale bar = 350µm (left panels) and 50 µm (right panels). (F) Representative photographs of subcutaneous xenografts from mice fed matched high (158 µm) and low Cu (4 µM) diet. Scale bar =1 cm. (G) Volume of subcutaneous tumors measured during 6 weeks of tumor growth. (H) Weight of subcutaneous xenografts at the time of collection (7 weeks after injection). (I) Total tumor Cu levels in xenograft tumors measured by ICP-MS. (J) Allocation of Cu to CuCOX measured by SEC-ICP-MS. (K) Schematic representation of protocol to adpart RCC cell line to chronic high Cu levels in the media. (L) Pathways identified by GSEA enriched in Cu*^Hi^* vs. Cu*^Lo^* 786-O cells. Means ±SEM shown; P values from two tailed t-test. See also Figure S1, Table S1 and Table S2A.

**Figure S2.**
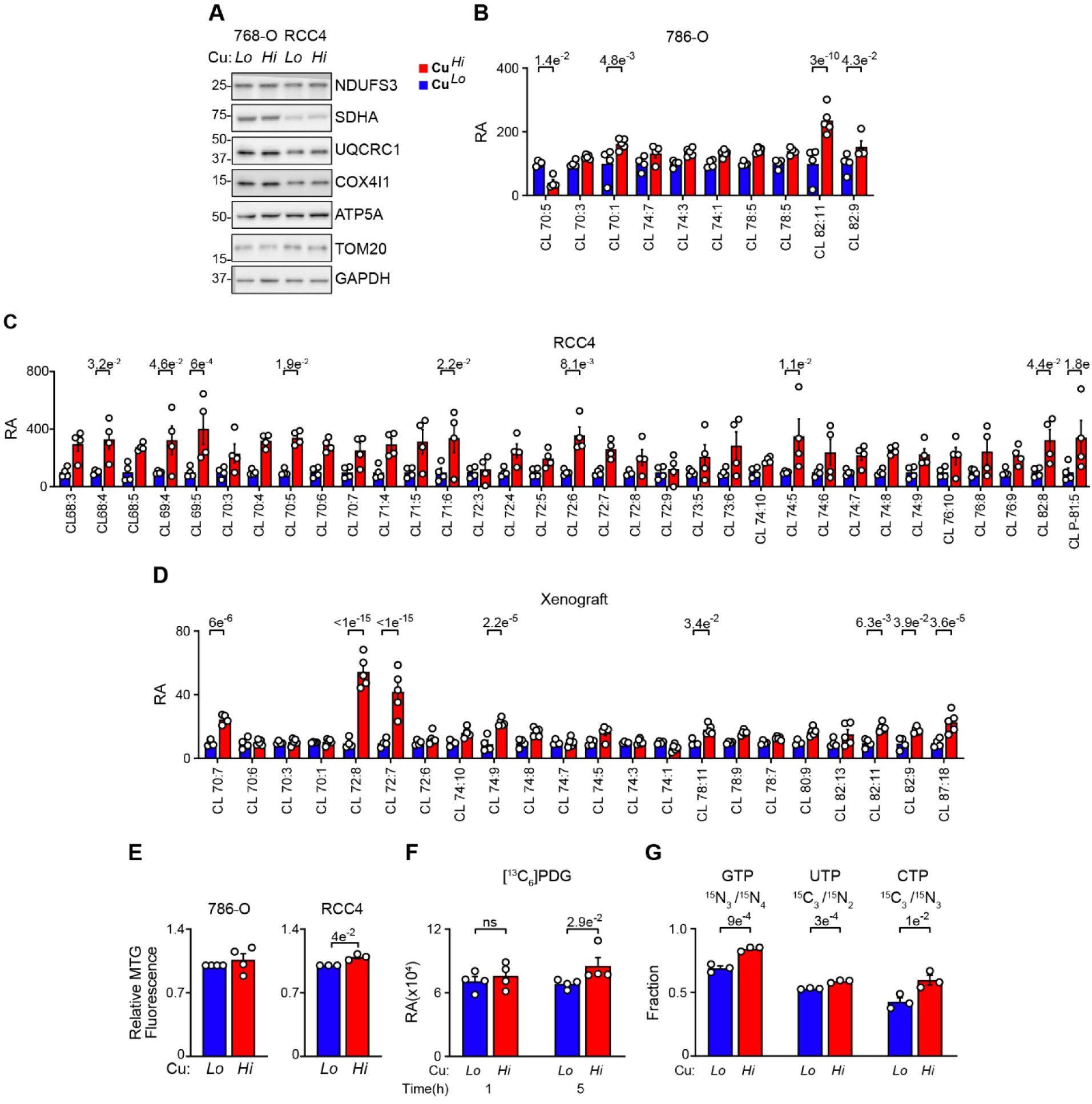
**related to Figure 2. Effects of Cu on metabolism** (A) Western blot of the subunits of the respiratory complexes in whole cell lysates. (B&C) Speciation of cardiolipins affected by Cu treatment in 786-O and RCC4 cells. (D) Speciation of cardiolipins in xenograft tumors formed in mice fed low and high Cu diet. Data is square root transformed. (F) MitoTracker Green fluorescence in Cu*^Lo^* and Cu*^Hi^* cells. (G) Cu effects on labeling of 6PDG, intermediate of oxidative branch of PPP, after indicated incubations with [^13^C_6_]Glc. (H) Cu effects on labeling GTP, UTP and CTP from [^13^C_5_,^15^N_2_]Gln after 24 h incubation. (I) Means ±SEM shown; P values in A, B, G, I calculated from two tailed t-test. P-value in B-D calculated by two-way ANOVA. See also Tables S2A, S2B and S3.

**Figure S3.**
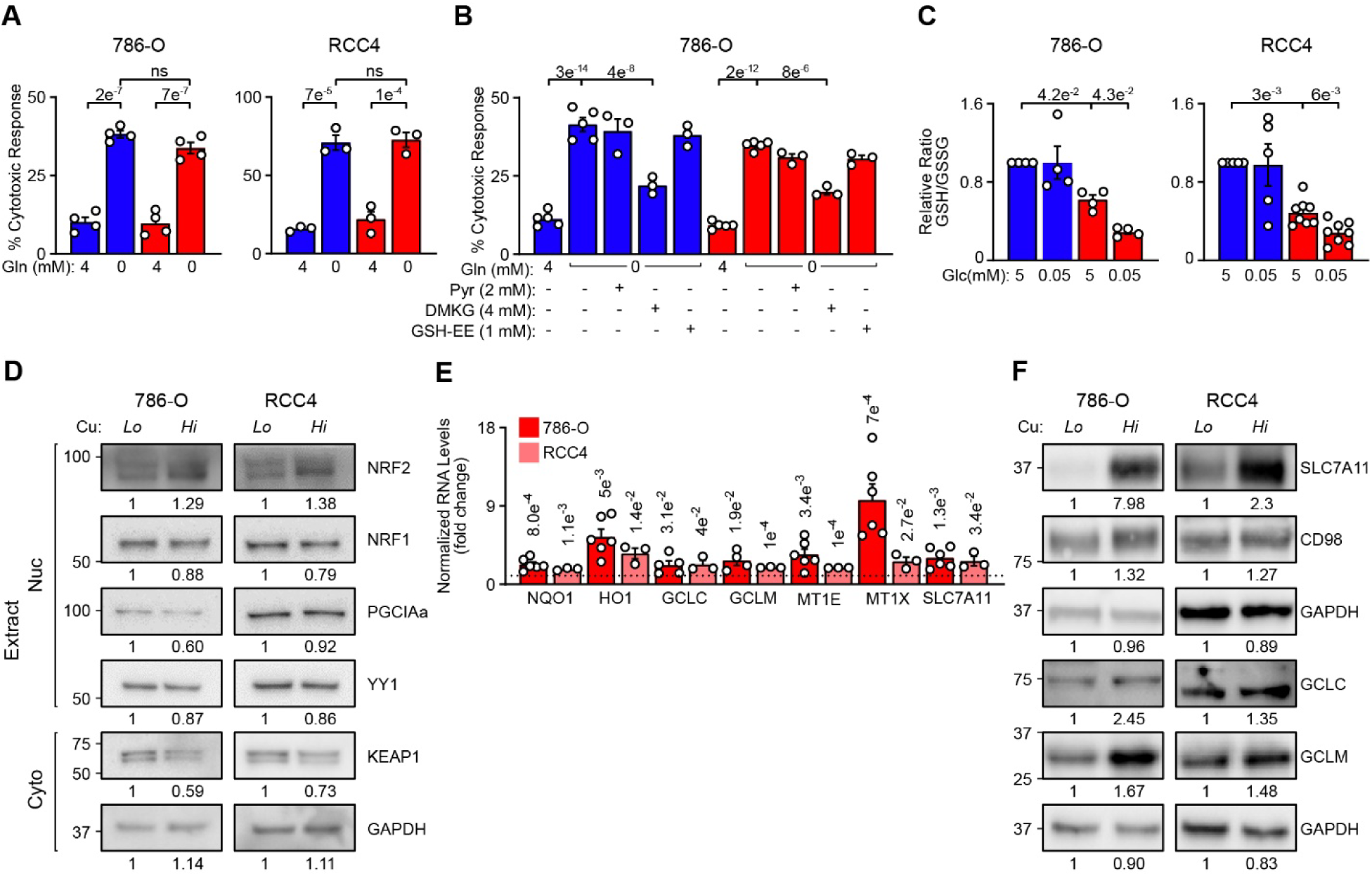
**related to Figure 3. Cu induces NRF2 -dependent activation of glutathione biosynthesis.** (A) Cytotoxic effect of glutamine starvation in Cu*^Lo^* and Cu*^Hi^* 786-O and RCC4 cells. (B) Partial rescue of cell death with DMKG but not pyruvate or GSH-EE in response to glutamine starvation in Cu*^Lo^* and Cu*^Hi^* 786-O cells. (C) Effects of glucose starvation on GSH/GSSG ratios in Cu*^Lo^* and Cu*^Hi^* cells. (D) Western blot of nuclear (Nuc) and cytosolic (Cyto) extracts probed for indicated proteins in 786-O and RCC4 Cu*^Lo^* and Cu*^Hi^*cells. (E) qRT-PCR of the indicated NRF2 targets in 786-O and RCC4 cells. Data presented as ratios of measurements in Cu*^Hi^* cells to that in Cu*^Lo^* cells. P values from two tailed t-test. (F) Representative western blot of the indicated proteins required for glutathione biosynthesis in Cu*^Lo^* and Cu*^Hi^* 786-O and RCC4 cells. Means ±SEM shown; In A-C, P values calculated from one way ANOVA with Holm-Sidak post-test.

**Figure S4.**
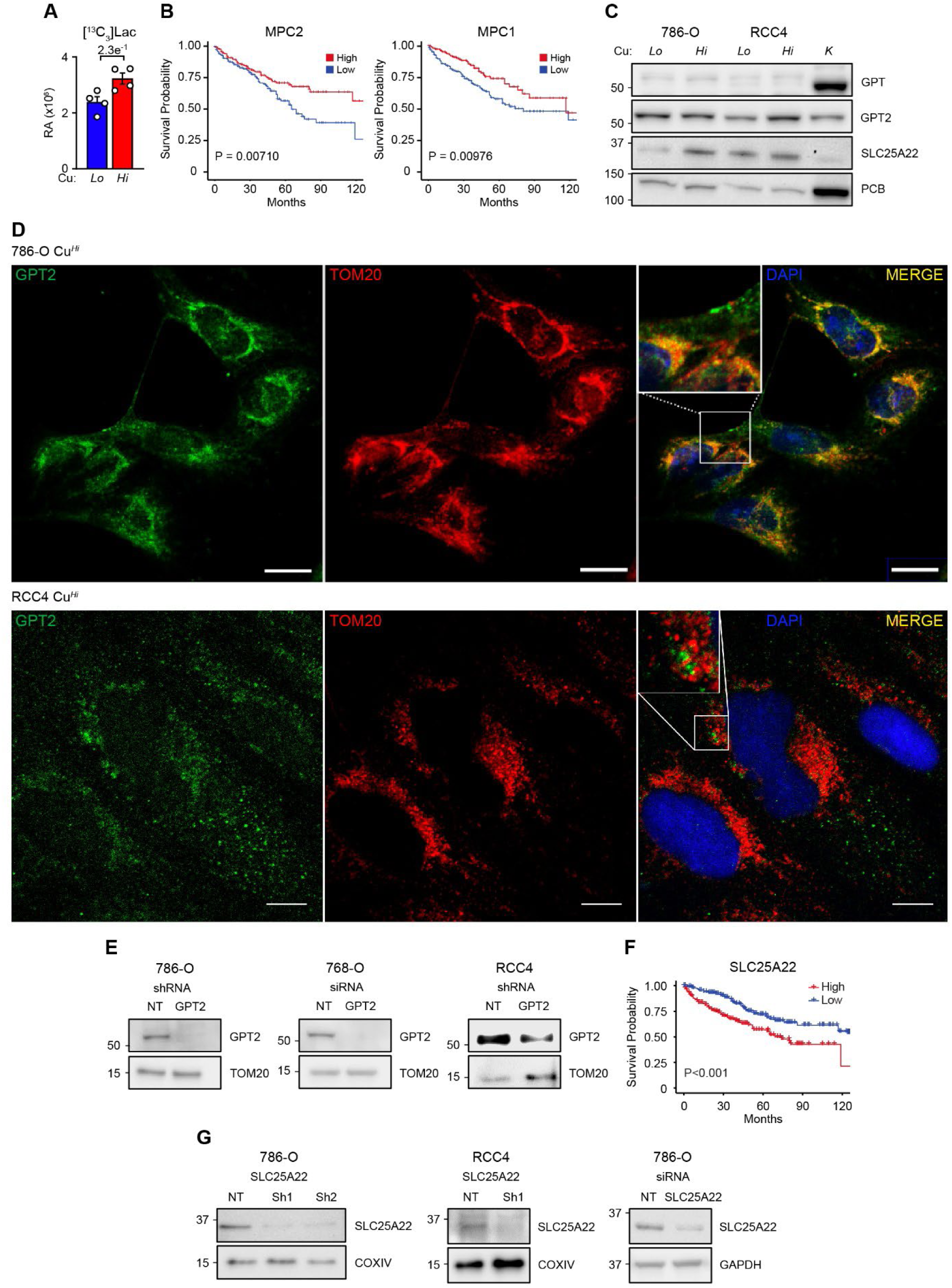
**related to Figure 4. Glucose and glutamine metabolism in Cu*^Hi^* and Cu*^Lo^* cells.** (A) Relative abundance (RA) of [^13^C_3_]-lactate (Lac) in Cu*^Lo^* and Cu*^Hi^* 786-O cells treated with [^13^C_6_]Glc (5 h). Means ±SEM shown; P value from two tailed t-test. (B) Kaplan Meier curves show predicted survival based on the levels of MPC1 and MPC2 in TCGA KIRC database. (C) Western blot showing expression of GPT1, GPT2 and SLC25A22 in the total cellular lysates in Cu*^Lo^* and Cu*^Hi^* cells. K-normal human kidney extract. (D) Immunofluorescence for GPT2 shows mitochondrial and cytosolic localization in Cu*^Hi^* 786-O and RCC4 cells. Scale bars = 20 µm for 786-O cells and 10 µm for RCC4 cells. (E) Western blot showing effective knockdowns of GPT2 in Cu*^Hi^* 786-O and RCC4 cells. (F) Kaplan Meier curves show predicted survival based on the levels of SLC25A22 in TCGA KIRC database. (G) Western blot showing effective knockdowns of SLC25A22 in Cu*^Hi^* 786-O and RCC4 cells.

**Figure S5.**
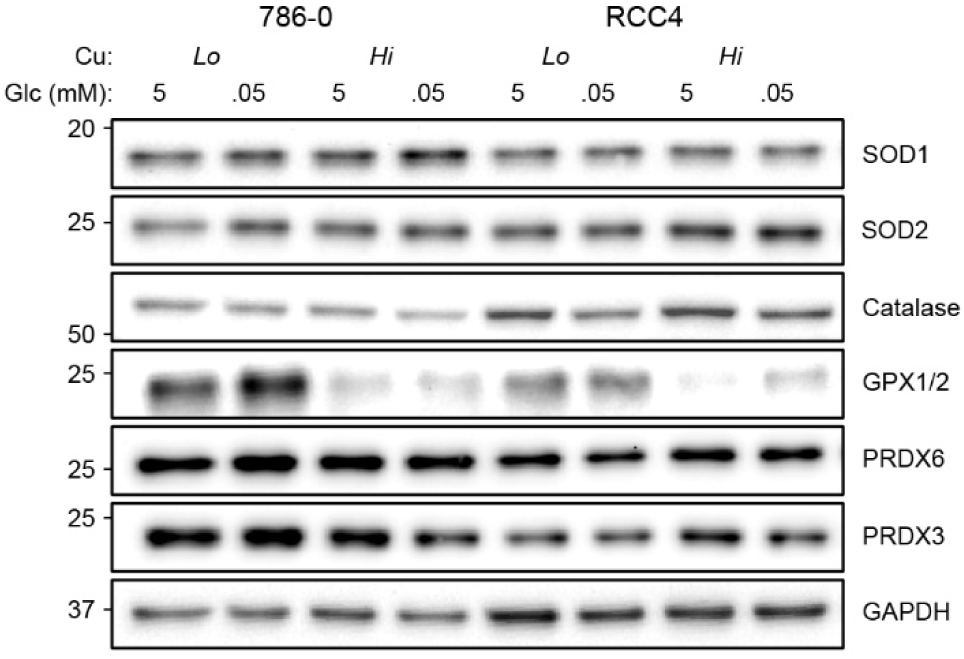
**related to Figure 5. Western blot showing expression of indicated enzymes generating and decomposing H_2_O_2_ in Cu*^Lo^* and Cu*^Hi^* 786-O and RCC4 cells at the indicated glucose concentrations.**

**Figure S6.**
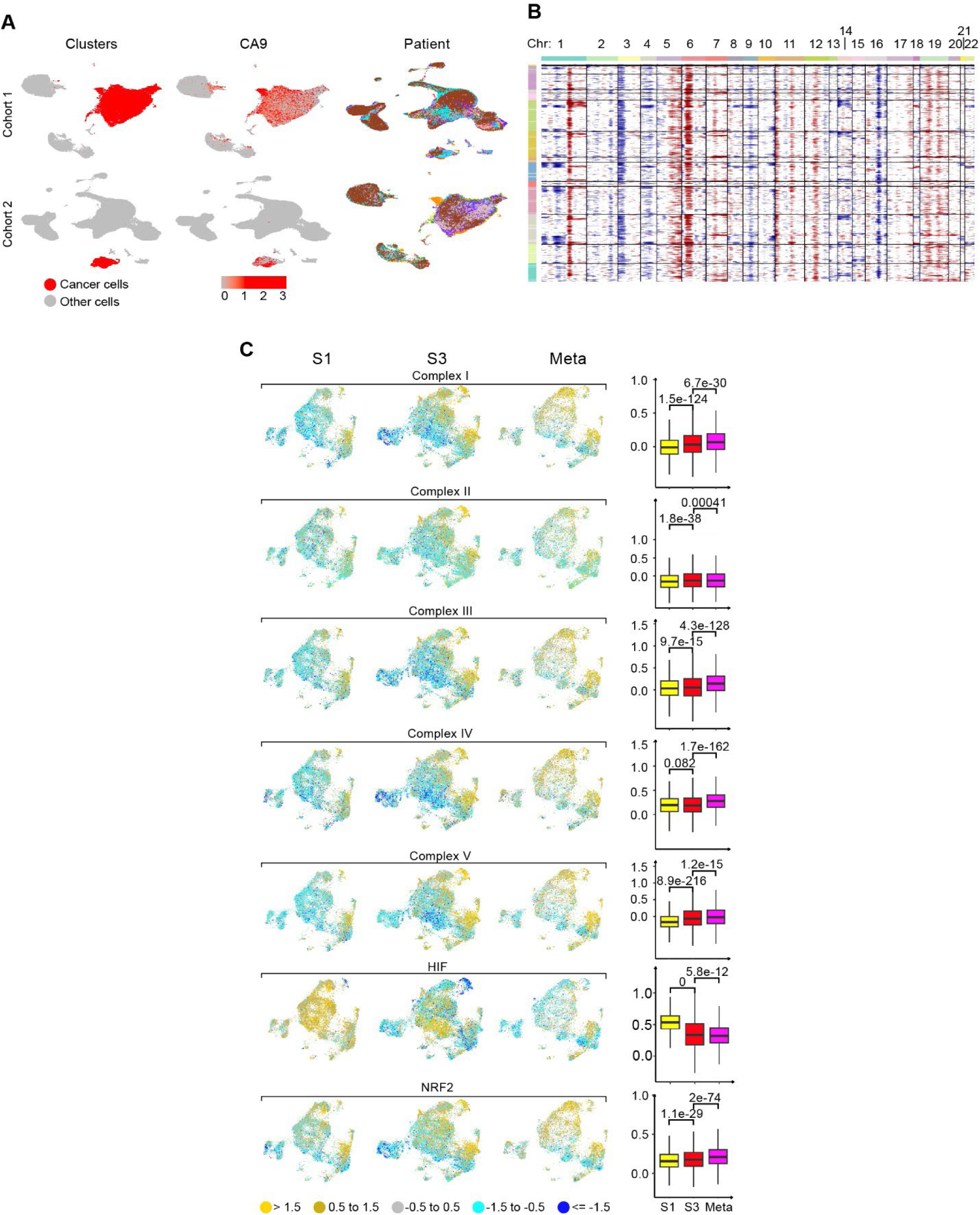
**related to Figure 6. Characterization of subpopulation of cancer cells identified by scRNA-seq.** (A) Identification of ccRCC cancer cells from two data sets of ccRCCs cohorts. Left: Identification of all cell subpopulations. Cancer cells are labeled red. Middle: Expression of HIF target carbonic anhydrase 9 (CA9) is used for the identification of cancer cells. Right: Mapping the data from the individual patients into subpopulation of cells. (B) Inferred copy number variants for the combined cohort of tumors. Top - chromosome number; left-subpopulations of cancer cells. (C) UMAPs visualizations (left) show overall increase in expression of genes encoding individual respiratory complexes during advancement of ccRCC, and NRF2 targets while reduction in HIF targets in more advanced ccRCCs. Box-whisker plots (right) show the distributions of module scores for each pathway signature across all putative cancer cells. Each column shows expression of the indicated gene set using the Seurat module score defined relative to a control gene set.

**Figure S7.**
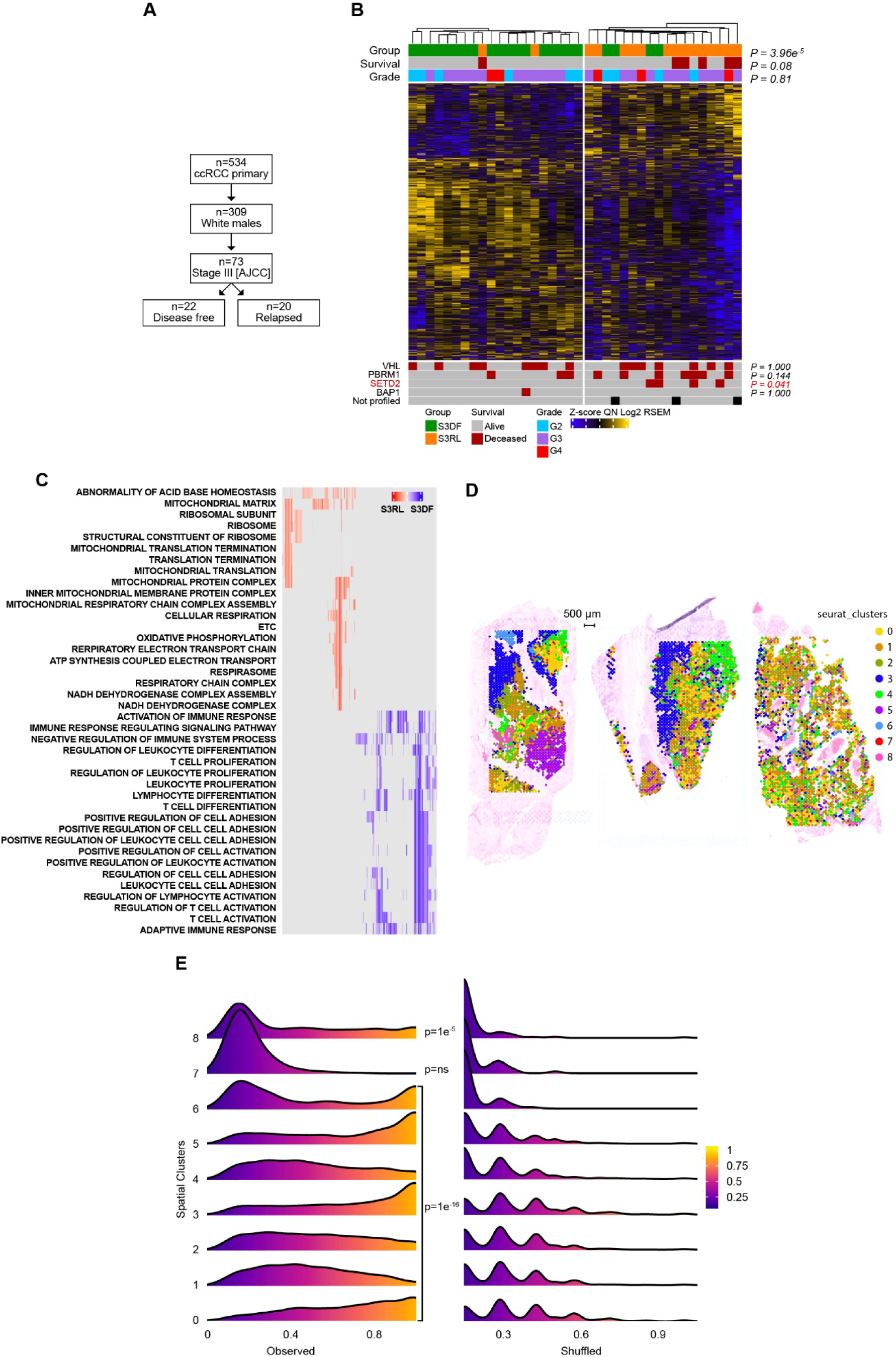
**related to Figure 7. Bulk RNA and spatial transcriptomics analysis of ccRCC.** (A) Identification of the cohort of white males with stage 3 ccRCC who remained disease free (S3DF) or relapsed (S3RL) within 2 years after surgery from TCGA KIRC Firehose cohort. (B) Clustered heatmap of differentially expressed genes (t-test, P<0.05) stratifies the cohort into predominantly S3DF or S3RL groups. Fisher exact test was used to quantify the association. (C) Heatmap showing clustered genes in leading edge subsets of top 20 most enriched GSEA ontology gene sets significantly associated with genes differentially regulated in S3DF and S3RL. Pathways associated with mitochondrial activity, mitochondrial translation and electron transport chain are enriched in S3RL tumors, while pathways associated with immune and inflammatory responses are enriched in S3DF tumors. (D) Mapping of 9 spatial clusters into H&E section of three ccRCCs. (E) Histogram of localization of each spatial cluster in the observed (left) vs. randomly shuffled spots (right). Numbers close to 1 indicate that the neighbors of a spot belong to the same spatial cluster. P values from Kolmogorov-Smirnov test measure the difference between the observed vs. shuffled distribution of each cluster localization.

