## Supplementary material for "Copper drives remodeling of metabolic state and progression of clear cell renal cell carcinoma": Table S7

**STAR* METHODS**

**KEY RESOURCE TABLE**

| REAGENT or RESOURCE | SOURCE | IDENTIFIER |
| --- | --- | --- |
| Antibodies |  |  |
| ATP5A Mouse antibody [15H4C4] [WB 1:1,000] | Abcam | CAT#: ab14748; RRID:AB_301447 |
| Catalase (D4P7B) XP Rabbit mAb [WB 1:1,000] | Cell Signaling Technology | CAT#: 12980; RRID:AB_2798079 |
| CD31 rabbit antibody [IHC-1:25] | Abcam | CAT#: ab28364;  RRID:AB_726362 |
| CD98 Mouse Antibody (E-5) [WB 1:1,000] | Santa Cruz | CAT#: sc-376815; RRID:AB_2938854 |
| COX IV (COX4I1) Rabbit Antibody [WB 1:1,000] | Cell Signaling Technology | CAT#: 4844; RRID:AB_2085427 |
| COX17 rabbit antibody [WB 1:1,000] | My BioSource | CAT#: mbs2526660;  RRID:AB_3076222 |
| COX7A2L Rabbit Antibody [WB 1:1,000] | Proteintech | CAT#: 11416-1-AP; RRID:AB_2245402 |
| GAPDH mouse antibody [6C5] [WB 1:10,000] | Abcam | CAT#: ab8245;  RRID:AB_2107448 |
| GCLC mouse antibody (3H1) [WB 1:1,000] | Novus Biologicals | CAT#: H00002729-M01;  RRID:AB_2294387 |
| GCLM rabbit antibody [EPR6667] [WB 1:1,000] | Abcam | CAT#: ab126704  RRID:AB_11127439 |
| GPT/ALT1 rabbit antibody [WB 1:500] | Proteintech | CAT#: 16897-1-AP;  RRID:AB_2230815 |
| GPT2 (G-7) mouse antibody [WB 1:1,000] | Santa Cruz | CAT#: sc-398383;  RRID:AB_2927429 |
| GPT2 rabbit antibody [IF 1:100] | Proteintech | CAT#: 16757-1-AP;  RRID:AB_2112098 |
| GPX1/2 (B-6) mouse antibody [WB 1:1,000] | Santa Cruz | CAT#: sc-133160;  RRID: sc-133160 |
| KEAP1 (D6B12) Rabbit mAb [WB 1:1,000] | Cell Signaling Technology | CAT#: 8047; RRID:AB_10860776 |
| Ki-67 (30-9) rabbit CONFIRM antibody [IHC-used as supplied] | Roche Ventana | CAT#: 05278384001;  RRID:AB_2631262 |
| MT-CO1 Rabbit Antibody [WB 1:1,000] | Abcam | CAT#: ab203912; RRID:AB_2801537 |
| MT-CO2 Rabbit Antibody [WB 1:1,000] | Abbexa | CAT#: abx125706;  RRID:AB_3076223 |
| NDUFS3 Rabbit Antibody [WB 1:1,000] | Proteintech | CAT#: 15066-1-AP;  RRID:AB_2151109 |
| NRF1 (D9K6P) Rabbit mAb [WB 1:1,000] | Cell Signaling Technology | CAT#: 46743; RRID:AB_2732888 |
| NRF2 (D1Z9C) XP Rabbit mAb [WB 1:1,000] | Cell Signaling Technology | CAT#: 12721; RRID:AB_2715528 |
| PCB (D-9) mouse antibody | SantaCruz | CAT#: sc-365673;  RRID:AB_10842023 |
| PDH-E1α Mouse Antibody (D-6) [WB 1:1,000] | Santa Cruz | CAT#: sc-377092; RRID:AB_2716767 |
| PDK1 Rabbit Antibody [WB 1:1,000] | Cell Signaling Technology | CAT#: 3062;  RRID:AB_2236832 |
| PDP1/PPM2C rabbit antibody | Proteintech | CAT#: 21176-1-AP;  RRID:AB_2878824 |
| PGC-1α (3G6) Rabbit mAb [WB 1:1,000] | Cell Signaling Technology | CAT#: 2178; RRID:AB_823600 |
| Phospho-PDH E1 Alpha (Ser232) Polyclonal rabbit antibody [WB 1:1,000] | Proteintech | CAT#: 29582-1-AP; RRID:AB_2918327 |
| Phospho-PDH E1 Alpha (Ser300) Recombinant rabbit antibody [WB 1:1,000] | Proteintech | CAT#:80572-1-RR; RRID:AB_2918903 |
| PRDX3 rabbit antibody [WB 1:1,000] | Proteintech | CAT#: 10664-1-AP;  RRID:AB_2284207 |
| PRDX6 rabbit antibody [WB 1:1,000] | Proteintech | CAT#: 13585-1-AP;  RRID:AB_2168637 |
| Pyruvate Dehydrogenase E1-alpha subunit [p Ser293] Rabbit Antibody [WB 1:1,000] | Novus | CAT#: NB110-93479; RRID:AB_1237282 |
| SDHA (D6J9M) XP Rabbit antibody [WB 1:1,000] | Cell Signaling Technology | CAT#: 11998; RRID:AB_2750900 |
| SLC25A22 rabbit antibody [WB 1:1,000] | Proteintech | CAT#: 25402-1-AP;  RRID:AB_2880060 |
| SOD1 Rabbit Antibody [WB 1:1,000] | Cell Signaling Technology | CAT#: 2770; RRID:AB_2302392 |
| SOD2 Rabbit Polyclonal antibody [WB 1:1,000] | Proteintech | CAT#: 24127-1-AP; RRID:AB_2879437 |
| TOM20 (F-10) mouse antibody [WB 1:1000 and IF 1:2000] | Santa Cruz | CAT#: sc-17764;  RRID:AB_628381 |
| TOM22 (E3F4M) Rabbit mAb [WB 1:1,000] | Cell Signaling Technology | CAT#: 90704S;  RRID:AB_3076225 |
| UQCRC1 rabbit antibody [WB 1:1,000] | My BioSource | CAT#: mbs9415267;  RRID:AB_3076226 |
| xCT/SLC7A11 (D2M7A) Rabbit mAb [WB 1:1,000] | Cell Signaling Technology | CAT#: 12691; RRID:AB_2687474 |
| YY1 Mouse Antibody (H-10) [WB 1:1,000] | Santa Cruz | CAT#: sc-7341; RRID:AB_2257497 |
| Goat anti-rabbit IgG, HRP-linked Antibody [WB 1:5,000] | Cell Signaling Technology | CAT#: 7074; RRID:AB_2099233 |
| Horse anti-mouse IgG, HRP-linked Antibody [WB 1:5,000] | Cell Signaling Technology | CAT#: 7076; RRID:AB_330924 |
| Goat anti-Mouse IgG (H+L) Secondary Antibody, Alexa Fluor 555 conjugate [IF 1:1,000] | Thermo Fisher Scientific | CAT#: A-21422; RRID:AB_2535844 |
| Goat anti-Rabbit IgG (H+L) Cross-Adsorbed Secondary Antibody, Alexa Fluor 488 [IF 1:1,000] | Thermo Fisher Scientific | CAT#: A-11008; RRID:AB_143165 |
| Biological Samples |  |  |
| Human ccRCCs, frozen | University of Cincinnati Cancer Center Biospecimen Shared Resource | https://med.uc.edu/institutes/cancer/shared-resources/biospecimen-shared-resource |
| Chemicals, Peptides, and Recombinant Proteins |  |  |
| Copper (II) Sulfate Pentahydrate | Thermo Scientific | CAT#: AC423615000;  CAS#: 7758-99-8 |
| Buthionine Sulfoximine (BSO) | Cayman | CAT#: S9728;  CAS#: 83730-53-4 |
| Oligomycin A | Cayman | CAT#: 11342  CAS#: 579-13-5 |
| FCCP | Cayman | CAT#: 15218;  CAS#: 370-86-5 |
| Rotenone | Sigma | CAT#: R8875;  CAS#: 83-79-4 |
| Antimycin A | Sigma | CAT#: A8674;  CAS#: 1397-94-0 |
| Hoechst 33342 | Thermo Scientific | CAT#: 62249;  CAS#: 23491-45-4 |
| U-13C6 D-Glucose | Cambridge Isotope Laboratories | CAT#: CLM-1396;  CAS#: 110187-42-3 |
| 13C5, 15N2 L-Glutamine | Cambridge Isotope Laboratories | CAT#: CNLM-1275-H;  CAS#: 285978-14-5 |
| Sodium 2,3-13C2 Pyruvate | Cambridge Isotope Laboratories | CAT#: CLM-3507;  CAS#: 89196-78-1 |
| Glucose | Gibco | CAT#: A24940-01;  CAS#: 50-99-7 |
| L-Glutamine | Sigma | CAT#: G7513;  CAS#: 56-85-9 |
| Sodium Pyruvate | Sigma | CAT#: S8636;  CAS#: 113-24-6 |
| Seahorse XF 1.0 M Glucose Solution | Agilent | CAT#: 103577-100 |
| Seahorse XF 200 mM glutamine solution | Agilent | CAT#: 103579-100; |
| Seahorse XF 100 mM pyruvate solution | Agilent | CAT#: 103578-100; |
| Dimethyl 2-oxoglutarate (DMKG) | Sigma | CAT#: 349631;  CAS#: 13192-04-6 |
| Glutathione ethyl ester (GSH-EE) | Cayman | CAT#: 14953;  CAS#: 92614-59-0 |
| UK5099 | Sigma | CAT#: 56396-35-1;  CAS#: 56396-35-1 |
| GNE-140 | Sigma | CAT#: SML2580;  CAS#: 2003234-63-5 |
| L-Cycloserine | MyBioSource | CAT#: MBS575566;  CAS#: 339-72-0 |
| L-Alanine | Sigma | CAT#: 05129;  CAS#: 56-41-7 |
| 3,3'-Diaminobenzidine | Sigma | CAT#: D12384  CAS#: 91-95-2 |
| Cytochrome c from bovine heart | Sigma | CAT#: C3131  CAS#: 9007-43-6 |
| Catalase-polyethylene glycol (PEG-Catalase) | Sigma | CAT#: C4963 |
| Catalase-polyethylene glycol (PEG-Catalase) | Nanocs | CAT#: CTS-PEG-1 |
| Ammonium tetrathiomolybdate (TTM) | Sigma | CAT#: 323446;  CAS: 15060-55-6 |
| DAPI (4',6-Diamidino-2-Phenylindole, Dilactate) | Invitrogen | CAT#: D3571  CAS#: 28718-91-4 |
| Critical commercial assays |  |  |
| GSH Glo Glutathione Assay | Promega | CAT#: V6911 |
| Propidium Iodide/FITC Annexin V Kit Dead Cell Apoptosis Kit | Invitrogen | CAT#: V13242 |
| CyQuant NF Proliferation Assay | Invitrogen | CAT#: C35006 |
| MitoSOX Red | Invitrogen | CAT#: M36008 |
| MitoTracker Green FM | Invitrogen | CAT#: M7514 |
| CM-H2DCFDA | Invitrogen | CAT#: C6827 |
| Deposited Data |  |  |
| Raw RNA sequencing from 786-O cell line | This paper | GEO GSE250028; https://www.ncbi.nlm.nih.gov/geo/query/acc.cgi?acc=GSE250028); |
| Raw sequencing data from spatial transcriptomics | This paper | GEO GSE250163; https://www.ncbi.nlm.nih.gov/geo/query/acc.cgi?acc=GSE250163 |
| Codes | This paper | https://github.com/mjarek66git/ccRCC |
| Experimental Models: Cell Lines |  |  |
| 786-O | ATCC | CAT#: CRL-1932; RRID:CVCL_1051 |
| RCC4 |  | RRID:CVCL_0498 |
| Experimental Models: Organisms/Strains |  |  |
| Mouse: NU/J (Athymic nude) | The Jackson Laboratory | Strain #:002019  RRID:IMSR_JAX:002019 |
| Mouse: NOD.Cg-*Prkdc^scid^ Il2rg^tm1Wjl^*/SzJ (NSG) | The Jackson Laboratory | Strain #:005557  RRID:IMSR_JAX:005557 |
| Oligonucleotides |  |  |
| siRNA: ON-TARGET plus Non-targeting Control pool | Dharmacon | CAT#: D-001810-10-20 |
| siRNA: ON-TARGET plus human GPT2 SMARTpool | Dharmacon | CAT#: L-004173-01-0005 |
| siRNA: ON-TARGET plus human SLC25A22 SMARTpool | Dharmacon | CAT#: L-007482-01-0005 |
| Primer Set: NQO1  Forward: GGATGAGACACCACTGTATTT  Reverse: CTCCTCATCCTGTACCTT | This paper | N/A |
| Primer Set: HO1  Forward: GGGCCAGCAACAAAGTGCAAGATT  Reverse: TCGCCACCAGAAAGCTGAGTGTAA | This paper | N/A |
| Primer Set: GCLC  Forward: TGCCCAGAGTTACTTGGATCAGCA  Reverse: AGAGGCATGGTACTGTAGCCAGTT | This paper | N/A |
| Primer Set: GCLM  Forward: CTGCTGTGTGATGCCACCAGATTT  Reverse: GTGCGCTTGAATGTCAGGAATGCT | This paper | N/A |
| Primer Set: MT1E  Forward: CCCTTTGCTCGAAATGGA  Reverse: AACAGCAGCTCTTCTTGC | This paper | N/A |
| Primer Set: MT1X  Forward: TTTCCTCTTGATCGGGAACTC  Reverse: GGCACAGGAGCCAACAG | This paper | N/A |
| Primer Set: SLC7A11  Forward: GGCATTTGGACGCTACAT  Reverse: CACTACAGTTATGCCCACAG | This paper | N/A |
| Recombinant DNA |  |  |
| GPT2 shRNA 1 | Dharmacon | CAT#: RHS3979-201764341;  TRCN0000035024 |
| SLC25A22 shRNA 1 | Dharmacon | CAT#: RHS3979-200798849;  TRCN0000044569 |
| SLC25A22 shRNA 2 | Dharmacon | CAT#: RHS3979-200798847;  TRCN0000044568 |
| Software and Algorithms |  |  |
| Image J | (Schneider et al, 2013) | https://imagej.nih.gov/ |
| Image Studio Lite | Li-Cor | https://www.licor.com/bio/image-studio-lite/download |
| SigmaPlot v14/v15 | Inpixon | https://systatsoftware.com/sigmaplot/ |
| GraphPad Prism | GraphPad | https://www.graphpad.com/features |
| Seahorse Wave Desktop Software | Agilent | https://www.agilent.com/en/product/cell-analysis/real-time-cell-metabolic-analysis/xf-software/seahorse-wave-desktop-software-740897 |
| Other |  |  |
| Teklad copper deficient diet | Envigo | TD.80388 |
| Teklad 10ppm copper diet | Envigo | TD.220421 |
| HyClone DMEM/F12 1:1 Media | Cytiva | CAT#: SH30023.02 |
| DMEM, no glucose, no glutamine, no phenol red | Gibco | CAT#: A1443001 |
| DMEM/F12 without L-Glutamine, Hepes, or Glucose | Biowest | CAT#: L0091 |
| Seahorse XF DMEM medium | Agilent | CAT#: 103575-100 |
| Matrigel Basement Membrane Matrix, LDEV-free | Corning | CAT#: 354234 |
| TSKgel QC-PAK GFC 300 column (7.8x150 mm, 5µm) | Sigma | CAT#: 816049 |
| Kinetex 1.7 µm C18 100 Å, LC Column 150 x 2.1 mm | Phenomenex | CAT#: 00F-4475-AN |
| Accucore C30 column: 2.6 μm, 2.1x150 mm | Thermo Scientific | CAT#: 27826-152130 |

**CONTACT FOR REAGENTS AND RESOURCES SHARING:**
